# Pre-T cell receptor Self-MHC Sampling Restricts Thymocyte Dedifferentiation

**DOI:** 10.1101/2022.04.28.489872

**Authors:** Jonathan S. Duke-Cohan, Aoi Akitsu, Robert J. Mallis, Cameron M. Messier, Patrick H. Lizotte, Wonmuk Hwang, Matthew J. Lang, Ellis L. Reinherz

## Abstract

Programming T lymphocytes to distinguish self from non-self is a vital, multi-step process arising in the thymus^1–4^. Signalling through the pre-T cell receptor (preTCR), a CD3-associated heterodimer comprising an invariant pTα chain and a clone-specific β chain, constitutes a critical early checkpoint in thymocyte development within the αβ T-cell lineage^5, 6^. Recent work demonstrates that preTCRs arrayed on double negative (DN) thymocytes, like αβ TCRs appearing on double positive (DP) thymocytes, ligate peptides bound to MHC molecules (pMHC) on thymic stroma but via a different molecular docking strategy^7–10^. Here we show the consequences of those distinctive interactions for thymocyte progression, using synchronized fetal thymic progenitor cultures differing in the presence or absence of pMHC on support stroma, determining single cell transcriptomes at key thymocyte developmental transitions. Although MHC negative stroma fosters αβ T lymphocyte differentiation, the absence of pMHC-preTCR interplay leads to deviant thymocyte transcriptional programming associated with de-differentiation. Highly proliferative DN and DP subsets with antecedent characteristics of T cell lymphoblastic and myeloid malignancies emerge. Thus, at least *in vitro*, beyond fostering β chain repertoire broadening for subsequent αβ TCR utilization, preTCR-pMHC interaction limits cellular plasticity to facilitate normal thymocyte differentiation and proliferation that, if absent, introduces significant developmental vulnerabilities.

---

The αβ T cell repertoire consists of many millions to billions of T lymphocytes, each expressing unique surface TCRs in a clonal manner^11–13^. These lymphocytes mediate precise recognition and elimination of aberrant host cells displaying “foreign” surface pMHC ligands consequent to infection or cellular transformation. In the thymus of jawed vertebrates during foetal, neonatal and juvenile life, the repertoire of clonotypic αβTCRs and their predecessor preTCRs is generated^6^. To this end, thymic progenitors originating from the bone marrow (and foetal liver *in utero*) proliferate during the early CD4^-^CD8^-^ double negative (DN1, DN2) stages and, under the influence of Notch at DN2, commit to the T cell lineage (Fig. 1a)^1^. Progression to the DN3a compartment (CD44^-^CD25^+^CD28^lo^) leads to further commitment to the αβT cell lineage, with recombination activating genes 1 and 2 (*Rag-1* and *Rag-2*) expression fostering TCRβ locus rearrangements that produce a recombined β chain expressed as a disulphide-linked heterodimer with the invariant pTα subunit^14^. In turn, pTα-β associates with the CD3 signalling subunits. Subsequently, upon preTCR signalling at the β selection checkpoint, the DN3b population (CD44^-^CD25^+^CD28^hi^) undergoes a critical program change to suppress Notch signalling, downregulate transcription of *Rag-1,-2* and *Ptcra* genes, increase cell cycling, and mediate allelic exclusion at the TCRβ locus enforcing expression of only one TCRβ chain per cell^6^. In turn, those thymocytes transition into the DN4 (CD44^-^CD25^-^) and then immature CD8 single positive (ISP) compartments^15^. Upon further progression to the double positive (CD4^+^CD8^+^; DP) stage, *Rag* genes are upregulated for a second time, permitting recombination and transcription at the TCRα locus and thereafter expression of the TCRαβ heterodimer^16–18^. To refine the αβ T cell repertoire, both positive and negative selection events ensue at this DP stage in the thymic cortex and continue into the maturing SP (CD4^+^CD8^-^ and CD4^-^CD8^+^) medullary compartment followed by their later export as peripheral T cells^19^.

**Fig. 1.**
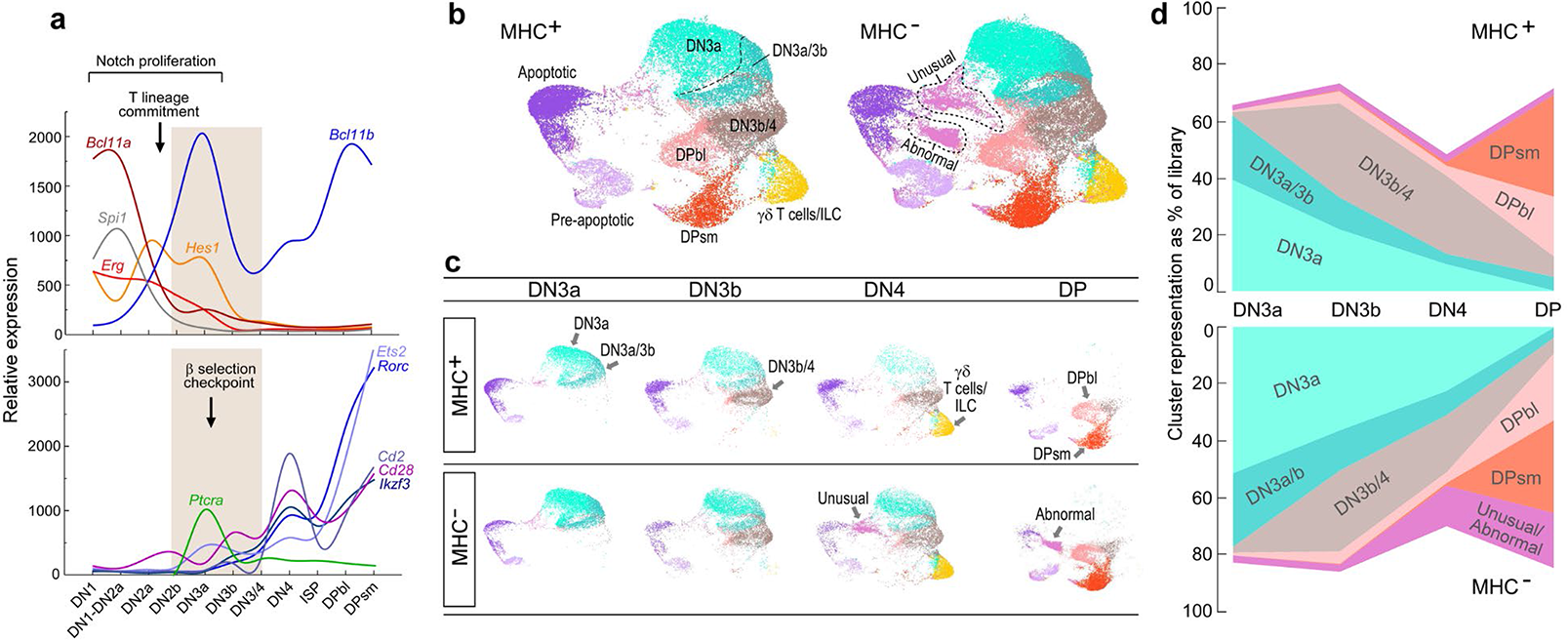
Developmental trajectories for thymocyte-like development on MHC^+^ or MHC^-^ supporting stroma. **a.** Schematic depicting representative gene transcript levels during key thymocyte developmental transitions (based on array data from the Immune Genome Project^25^). Upper panel: Early DN1-DN3 proliferation is driven by thymocyte Notch signalling, represented here by the *Erg* and *Hes1* transcripts. Myeloid development is suppressed during the DN2a to DN2b transition by downregulation of *Spi1* (coding for PU.1) and by the switch from *Bcl11a* to *Bcl11b* committing progenitors to the T lineage. Lower panel: following entry in to the DN3 stage and commitment to the αβ T cell lineage, preTCR with invariant pTα (pTCRα) is expressed. PreTCR signalling leads to downregulation of the Notch pathway, inhibition of TCR β locus recombination, downregulation of *Ptcra* and upregulation of the indicated transcripts. Curves have been smoothened between data points for each stage to aid tracking. **b.** UMAP projection of *k*-means clustering (*k* = 10) for DN3a, DN3b, DN4, and DP libraries for cells developing on either MHC^+^ or MHC^-^ stroma. All libraries are projected into the same space to permit direct comparison. Process for assignment of labels to each cluster is defined in the text. **c.** Cluster developmental trajectories of individual libraries and relationship of individual clusters to phenotypically characterized thymocyte subsets. Projection of the individual FACS-sorted libraries (labeled in large font) into the primary space allows initial assignment of clusters expressing distinct transcriptomes (labelled in small font) and permits identification and developmental staging of cluster differences between the MHC^+^ and MHC^-^ conditions. **d.** MHC^-^ thymocyte-like cells progress developmentally by phenotype from DN3a to the DP stage but with altered distribution on comparison with MHC^+^ cells. To focus only on the αβTCR lineage, ILC-γ/δ-like T cells and pre-apoptotic/apoptotic cells are excluded. For each library, the proportion of each defined developmental cluster is depicted. Cluster colours are consistent with those used in Figs. 1b and 1c. For each MHC^+^ library, cell numbers in parentheses: DN3a (6,970 cells), DN3b (7,711 cells), DN4 (7,337 cells), and DP (5,747) and likewise for the MHC^-^ libraries: DN3a (9,453 cells), DN3b (8,776 cells), DN4 (12,454 cells), and DP (13,011). P < 2.5 x 10^-7^ for difference between MHC^+^ and MHC^-^ stage distributions (Chi-square statistic).

PreTCR signalling was judged independent of ligand recognition at the DN3 stage consequent to several lines of prior investigation^20–23^. First, ablation of the TCR β chain variable domain that forms part of the interaction surface with pMHC in the αβTCR did not impact development through the DN3a to DN3b checkpoint. Second, in further support of ligand binding dispensability, a preTCR missing the extracellular domains of both the β chain and the pTα chain could drive development to the DP compartment. Third, in MHCI^-^MHCII^-^ double knockout mice, thymocyte progression was unimpaired through the DN3 stage to the DP stage with respect to both cell numbers and phenotypes.

Recent structural and biophysical data, however, reveal direct interactions between preTCRs and pMHC ligands that utilize a horizontal binding mode compatible with facile mechanosensing^8, 10, 24^. Moreover, functional assays demonstrate both restricted proliferation and repertoire development in the absence of stromal MHCI and MHCII molecules^7, 8^. Together these findings necessitate re-examination of the earlier results. Therefore, we have investigated whether and how the absence of ligand-dependent preTCR signalling impacts proper thymocyte development. Through single cell transcriptome (scRNA-Seq) and bulk RNA-Seq analyses our study reveals cellular aberrations at both DN4 and DP stages consistent with a vital role of preTCR-pMHC interactions in enforcing orderly thymocyte-like transcriptional programming.

## Early T-lineage differentiation

Utilizing an *in vitro* model of thymocyte differentiation, we seeded haematopoietic stem cells (HSC) from foetal liver of wild-type C57Bl/6 mice onto OP9-DL4 MHCI (MHC^+^) stromal support cells or the same cells rendered MHCI-negative by CRISPR/Cas9 targeting of *B2m* and *Tap2* genes (MHC^-^)^7^. Both stromata lack endogenous MHCII expression. Extensive use of this model demonstrates synchronized expansion and development through to the immature single-positive (ISP) and DP stage within the d8 to d13 window thus recapitulating embryonic development dominated by a highly proliferative blast and DP compartment. This is in contrast to postnatal thymic development where formation of medullary components leads to robust presence of mature SP populations^25^.

To examine the TCR β chain selection checkpoint at the DN3a to DN3b transition, 3.2 x 10^4^ HSC were seeded onto MHC^+^ or MHC^-^ stroma and developing thymocyte-like cells analysed at d9 (6.225 x 10^7^ on MHC^+^; 3.375 x 10^7^ on MHC^-^). For brevity, we refer to cells generated on MHC^+^ and MHC^-^ stroma with prefix MHC^+^ or MHC^-^, respectively. Cells were sorted by FACS into DN3a, DN3b, DN4 and DP populations (Extended Data Fig. 1) and processed for scRNA-Seq using the 10X Genomics Chromium system simultaneously preparing from each cell a library enriched for TCR α and β chain clonotype transcripts. To reduce dimensionality of the transcriptome information, all libraries were aggregated and projected into a single Uniform Manifold Approximation and Projection (UMAP) plane allowing direct comparison of clusters and inferred trajectory analysis incident to the FACS sorting by phenotype (Fig. 1b; Supplemental Information Files 2, 3). To objectively delineate the relation of each cluster to thymocyte developmental stage, the dominant markers of normal transition from the DN3a to DPsm stages were extracted as reference arrays (Extended Data Fig. 2) from the Immunological Genome Project (IGP) α/β T lineage database^26^ and applied to each cluster yielding a transcriptome reference trajectory that matched with relative cluster representation in each stage-specific library (Fig. 1c). Within the DN4 libraries, a population with an γ/δ T cell-like and innate lymphoid cell-like (γ/δ-ILC) transcriptome signature partitions due to lack of expression of CD44 and CD25 (Extended Data Fig. 3a) pointing to the developmental fidelity of this *in vitro* system.

For this analysis, the pro-apoptotic (Extended Data Fig. 3b) and apoptotic populations (transcripts dominantly of mitochondrial origin) are retained not only as topological markers but to highlight the possibility, given the absence of thymic reticuloendothelial cells removing damaged cells, that in this assay apoptosis may be a significant process even before negative selection events occurring at DP stages and beyond. The DN3a/3b cluster (Fig. 1c) shows early upregulation of *Ikzf3* and *Cd28*, markers of preTCR signalling (Fig.1a, Extended Data Fig.2a) and bridges the DN3a and DN3b libraries. Likewise, the DN3b/4 cluster is represented in the DN3b, DN4, and DP libraries indicating the increased resolution over phenotype provided by the transcriptional signature (Fig. 1c, Extended Data Fig. 2b). The DPbl population segregates away from the mature DPsm population based on strong representation of cell cycling-related transcripts (Extended Data Fig. 2d).

## Lack of MHC impacts preTCR signalling

Having established the cluster signature trajectory in normal developmental progression, development in the MHC^-^ state was examined (Fig. 1b, 1c). The MHC^+^ trajectory for the DN3a cluster shows a clear diminution with progression from the DN3a to the DN4 libraries (Fig. 1c, top row and Fig. 1d). For the MHC^-^ state, this progression is significantly less where >36% of the DN3b cells by phenotype, and >22% of the phenotypically DN4 cells, retain a DN3a-like transcriptome, contrasted to 22% and <10%, respectively, in the MHC^+^ condition (Fig. 1c, d). Nonetheless, there is phenotypic developmental progression in the absence of potential pMHC ligand binding to the preTCR. The DN3a to DN3b transition is marked by upregulation of a new transcriptional program highlighted by upregulation of *Ikzf3*, *Rorc*, *Cd2*, *Cd28* and downregulation of *Hes1*, *Erg* and *Ptcra* (Fig. 1a, Extended Data Fig. 2b). We applied this gene panel to a subset of the DN3b/4 cluster more strongly represented in the MHC^-^ DN4 library than in the control condition, highlighted as a “tail” moving back into the DN3a/3b cluster (Fig. 1b, Fig. 2a). Splitting the MHC^-^ DN4 cluster into 2 subclusters, one representing the main region overlapping in position with the MHC^+^ DN3b/DN4 cluster and the other, the tail, showed clear differences. The latter, despite being phenotypically DN4, had neither upregulated *Ikzf3*, *Rorc* or *Cd2* nor downregulated *Hes1* and *Erg* as observed in the MHC^+^ cluster and, further, had not robustly upregulated *Trbv* transcription (Fig. 2b, Extended Data Fig. 2b). Collectively, these observations are consistent with a differentiation trajectory that bypasses the β selection checkpoint. The main MHC^-^ DN3b/4 cluster shows an intermediate expression between the MHC^+^ DN3b/4 cluster and the tail suggesting that elements of the aberrant transcriptional regulation observed in the tail subcluster extend to the main subcluster.

**Fig. 2.**
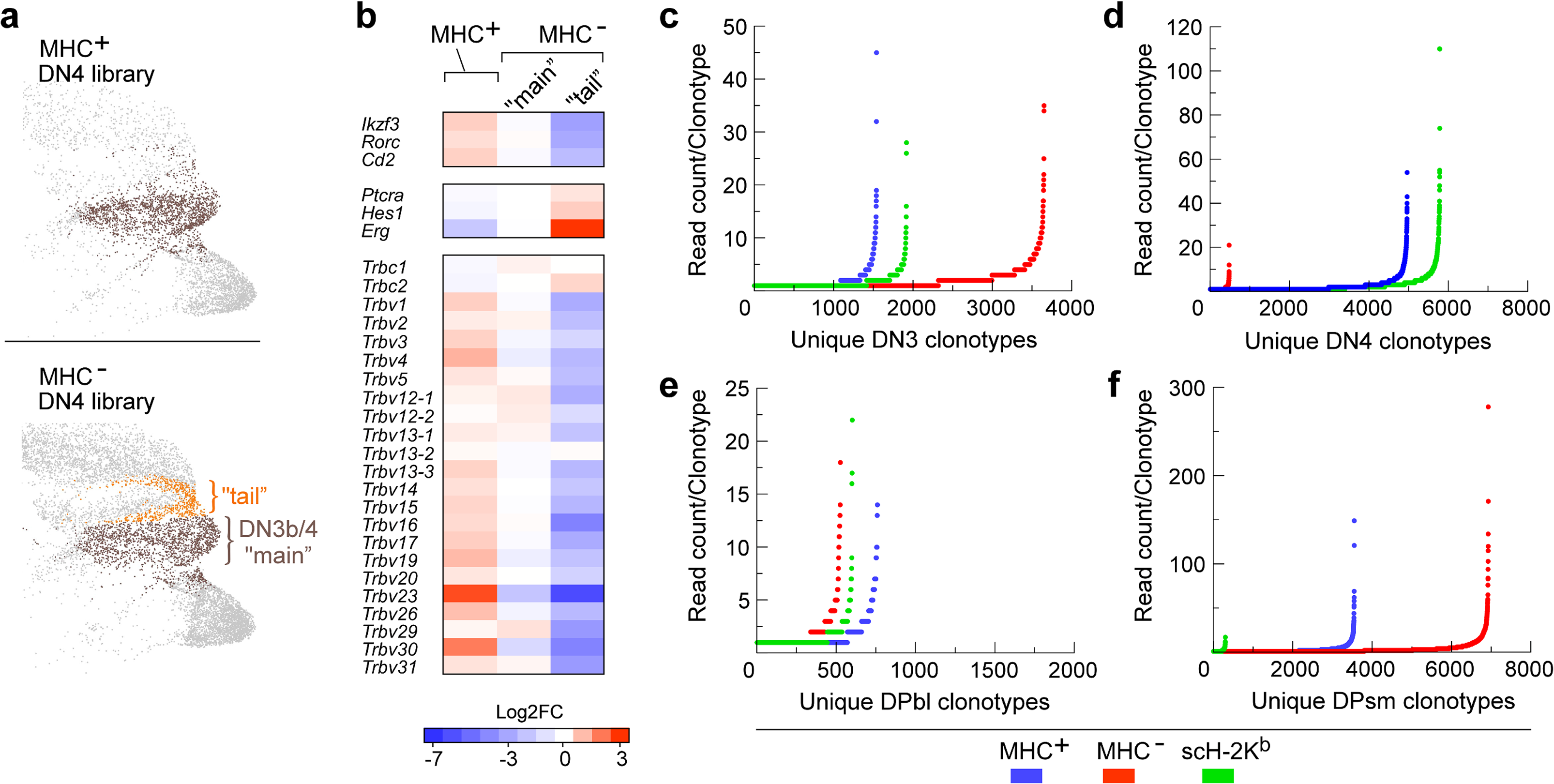
Uncoupling of the transcriptome and repertoire from phenotype in thymocyte-like cells developing on MHC^-^ stroma. **a.** The DN3b/4 cluster in the MHC^-^ DN4 library harbours a population with characteristics of cells not having passed through the preTCR signalling checkpoint. The MHC^-^ DN3b/4 cluster in the DN4 library is split into two subclusters, one corresponding to the DN3b/4 cluster in the MHC^+^ DN4 library (“main”, brown) and one corresponding to a set poorly represented in the MHC^+^ DN4 library (“tail”, orange). **b.** Transcript expression in MHC^+^ and MHC^-^ DN3b/4 cells. Transcripts well-expressed and marking the transition to DN3b/4 cells (*Ikzf3*, *Rorc*, *Cd2*; Suppl. Fig. 3) remain low in the MHC^-^ “tail” subcluster, transcripts expected to be downregulated remain high (Fig. 1a), and robust TCR β chain upregulation is not observed (Extended Data. Fig. 2b). **c-f.** Stage-specific analysis of β chain clonotype representation/ 10,000 cells in d9 MHC^+^, MHC^-^, and scH-2K^b^ OP9-DL4 thymocyte-like development cultures. Representative of 6 experiments examining MHC^+^ (n = 5), MHC^-^ (n = 6), scH-2K^b^ (n = 3).

## Reduced DN4 β clonotypic diversity

The importance of preTCR interacting with MHC for appropriate developmental regulation of *Trbv* transcription and repertoire diversity at the DN4 stage was examined further by β chain clonotype analysis of the developing subpopulations using targeted RNA-Seq. Wild-type HSC were seeded onto MHC^+^ OP9-DL4 stromal cells, onto MHC^-^ OP9-DL4 stromal cells or onto the same MHC^-^ cells transfected to re-express MHC class I as a single chain VSV8 peptide/β2m/H-2K^b^ (scH-2K^b^). The scH-2K^b^ derivative expresses multiple copies of a single pMHC thus maintaining the potential for the horizontal binding mode to the preTCR but presenting a homogenous peptide, RGYVYQGL, derived from amino acids 52–59 of vesicular stomatitis virus nucleoprotein^7^. After 9 days, cell proliferation was uniformly better on the MHC^+^ stromal cells than on either the MHC^-^ or scH-2K^b^ support stroma (Extended Data Fig. 4a, b). Cells from each support stroma culture were sorted into phenotypically defined DN3, DN4, DPbl and DPsm populations (Supplemental Information File 3) and *Trbv* clonotypes of 10^4^ cells for each stage and condition identified by targeted RNA-Seq.

TCR β clonotype diversity is relatively high at the DN3 stage for cells developing on all variants of the OP9-DL4 support stroma used here (Fig. 2c). The DN4 compartment, however, reveals consistently contracted repertoire diversity only on the MHC^-^ support stroma (Fig. 2d, Extended Data Fig. 4c). Further, up to 70% of the clonotypes developing in the MHC^-^ DN4 population were found at <7.5% levels in the MHC^+^ and scH-2K^b^ populations (Extended Data Fig. 4d) suggesting these clonotypes may represent a restricted population of TCR responding to non-classical MHC or MHC-unrelated structures on the stromal surface. Conversely, ∼92% of the clonotypes expressed on cells developing on the MHC^+^ and scH-2K^b^ stroma, where preTCR-pMHC interaction can occur, are absent in the MHC^-^ cultures. The limited clonotype representation in the MHC^-^ developing DN4 population is not a consequence of restricted cell proliferation since clonotype diversity of DN4 cells developing on scH-2K^b^ stroma is as rich as that of the cells developing in the MHC^+^ condition (Fig. 2d, Extended Data Fig. 4c) despite similar cell representation of all 3 DN4 cell populations (10^4^ cells analysed/sample). The characteristics of cells developing on MHC^-^ stroma or scH-2K^b^ stroma both diverge from those on the MHC^+^ stroma during the DP stage (Fig. 2e, f; Extended Data Fig. 4c). Cells developing on scH-2K^b^ stroma reveal a contraction of β repertoire diversity, likely linked to limited positive selection afforded by a single peptide (i.e., VSV8) on scH-2K^b^ stroma. Of note, the N15β clonotype with known specificity for VSV8 peptide presented by H-2K^b^ appears in the top 20 clonotypes coming through at the DPsm stage on the scH-2K^b^ stroma (Extended Data Table 1). On the other hand, the MHC^-^ developing cells recover diversity at the DPbl and DPsm stages, often overshooting that of cells on the MHC^+^ stroma (Fig.2, Extended Data Fig. 4c) and indicate aberration of β chain transcriptional regulation when MHC-dependent preTCR signalling is circumvented. Continued Notch stimulation in the absence of preTCR signalling has already been demonstrated to permit differentiation through to the DP stages^27^.

## Origin of β diversity in MHCIa^-^ system

The development of TCR clonotypes in the MHC^-^ condition implies that thymocytes can develop and bypass the preTCR checkpoint in the absence of MHC, either via a ligandless mode or utilizing non-classical MHCI and MHCII molecules or additional ligands. A panel of non-classical MHCI (MHCIb) was compiled (Extended Data Table 2) and, following full transcriptome analysis of the OP9 MHC^+^ and OP9 MHC^-^ stromal cells (Extended Data Fig. 5a-c), expression of non-β2m dependent MHC were examined. Loss of CD1d surface expression, dependent upon β2m, was used as a functional validation marker of the CRISPR/Cas9 knockout in addition to loss of MHCI (Extended Data Fig. 5d) thus supporting our focus upon non-β2m dependent MHC. Of all potential candidates, transcriptome analysis identified only Raet-1d and Raet-1e as being significantly expressed at the transcriptome level with detectable surface protein expression but with no difference between MHC^+^ OP9-DL4 and the MHC^-^ OP9-DL4 variant (Extended Data Fig. 5e). Consequently, the origin of the “background” clonotypes comprising the repertoire at the DN4 and subsequent stages in the MHC^-^ condition, also found as a minor fraction of the total repertoires in the MHC^+^ and scH-2K^b^ conditions (Extended Data Fig. 4d), is uncertain but may involve non-MHC ligands or non-classical MHCIb ligands that may be surface-expressed without an absolute requirement for β2m or assembly of the peptide-loading complex.

## Without MHC, unusual DN4 cells develop

To further address the diminution in β chain representation at DN4 in the MHC^-^ condition, examination of the scRNA-Seq clustering is informative. Although the partitioning of the ILC-like and γ/δ T-like cells within the DN4 represents a β chain-low population (Fig.1c; Extended Data Fig. 3a), this is not the source of the difference as the representation of this cluster is similar between the MHC^+^ and MHC^-^ conditions. Apart from the ILC-like and γ/δ T-like cells, the DN3b/4 cluster is the only other significant representation in the thymocyte developmental path within the MHC^+^ DN4 subpopulation. These cells exhibit a robust upregulation of β chain transcript (264.6±74.3 log2-fold increase, median = 88.4; P < 0.0001) on transitioning from the DN3a/3b cluster (Extended Data Fig. 2b). In contrast, in the MHC^-^ DN4 population in addition to the DN3b/4 population, there remains a high representation of phenotypically defined DN4 cells with a DN3a-like transcriptome as well as an unusual population barely observed in the MHC^+^ condition (“unusual”; Fig. 1c, d). As described above, on comparison with the MHC^+^ DN4 library *Trbv* transcript expression, within the MHC^-^ DN4 library the DN3b/4 main population trends toward suppression (Fig. 2b), the DN3b/4 tail exhibits a significant suppression (7.65-fold down against MHC^+^ DN3b/4, P<0.0002; 4.48-fold down against the MHC^-^ DN3b/4 main cluster, P<0.0025), as do the DN3a-like cells (5.38-fold down against MHC^-^ DN3b/4; P<0.0001), and the MHC^-^ unusual DN4 cluster (Fig. 3a; Extended Data Fig.5f). The aggregated effect of all these phenomena may contribute to the low DN4 *Trbv* clonotype representation in the MHC^-^ condition (Fig. 2d, Extended Data Fig. 4c).

**Fig. 3.**
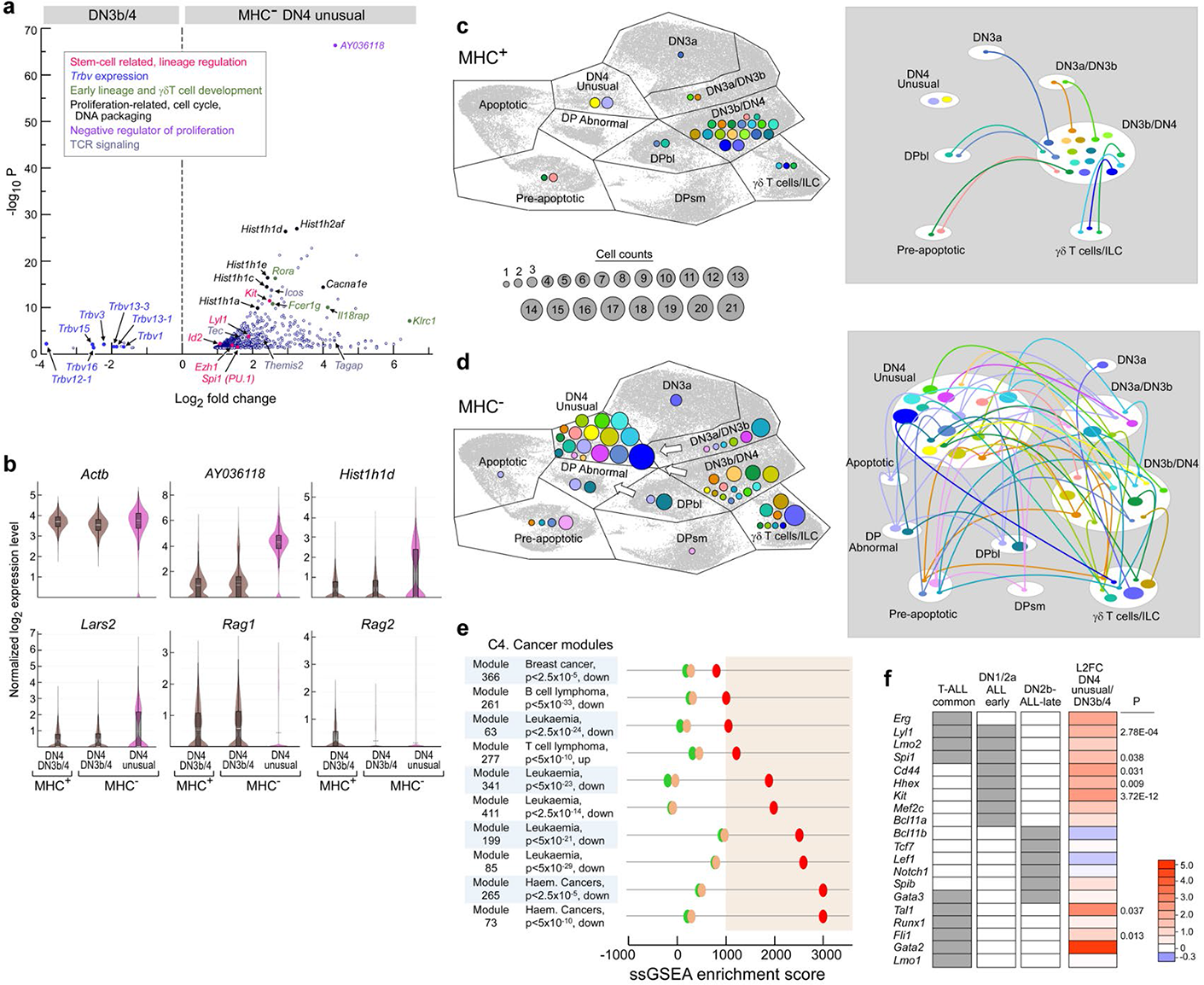
Single cell transcriptomics of the MHC^-^ DN4 unusual cluster reveal complex proliferative and lineage abnormalities. **a.** Volcano plot of significant transcript differences (P<0.05) between the DN4 unusual population and the DN3b/4 cluster in the same MHC^-^ DN4 library. Upregulated DN4 unusual transcripts are shown to the right of zero on the x-axis, downregulated to the left. Functional significance of the highlighted transcripts is listed in the inset box. An additional 14 *Trbv* transcripts were downregulated but did not meet the P<0.05 threshold. **b.** Violin plots depicting indicated log-normalised transcript levels selected for representation within most cells of the DN4 unusual cluster that are significantly different to representation with the DN3b/4 clusters within the MHC^+^, and MHC^-^ libraries, respectively. *Actb* transcript is included as a reference. Cluster colours are consistent with those depicted in Figs. 1 and 2. Box plot within cell distribution represents 1^st^ to 3^rd^ quartiles; where visible, dotted line within represents mean and solid line represents median. **c.** Cluster distribution of top 20 DN4 clonotypes developing on MHC^+^ stroma with expected developmental trajectory into DN3b/4 cluster. (left panel). Each colour represents a unique clonotype and circle diameter is proportional to the cell number expressing that specific β chain (see cell count scale common to Fig. 3c and Fig. 3d). Colour/clonotype specification is unique to the panel and bears no relation to colour use in Fig. 3d or Fig. 4c, d. Right panel depicts clonotype tracking from cluster to cluster with cluster/track colouring concordant with left panel. **d.** Distribution of top 20 DN4 clonotypes developing on MHC^-^ stroma similar to depiction in Fig. 3c. However, colour/clonotype specification is unique to the panel. Right panel depicts clonotype tracking from cluster to cluster and cluster/track colouring is concordant with left panel. **e.** ssGSEA scores for MHC^+^ DN3b/4 (green), MHC^-^ DN3b/4 (light orange), and MHC^-^ DN4 unusual (red) cells analysing module gene sets with co-ordinated gene regulation in tumours (MSigDb C4) and probability of association of specific tumors with up- or down-regulation of the specific module. ssGSEA scores greater than 1000 are considered strong, equal to zero as showing no correlation with module genes, and < 0 as inversely correlated. **f.** Comparison of the DN4 unusual cluster developing on MHC^-^ with the DN3b/4 cluster developing on DN4 MHC^+^ stroma (positive control) for transcripts reported as dysregulated in human T-ALL. Column1 grey boxes indicate regulatory transcripts found significantly over-expressed in human T-ALL in general. Column 2 grey box highlights regulatory genes of human T-ALL overexpressed in early thymic progenitors (ETP) characteristic of the pre-T lineage commitment checkpoint (labelled here DN1/DN2a ALL early). Column 3 highlights regulatory genes for transcript profiles of T-ALL consistent with a later DN2b stage (DN2b-ALL-late. Column 4 heatmap depicts log2-fold difference for the indicated transcript (scale to right); P indicates significance; if no value given then P>0.05).

The MHC^-^ unusual cluster (1776 cells; 14.3% of all DN4 cells), minimal in the MHC^+^ DN4 library (205 cells; 2.79% of all DN4 cells), displays a complex transcriptome. Unlike the DN3b/4 cells expected in the DN4 library, the unusual cluster cells have not consolidated the robust expression of β chains (Fig. 3a; Extended Data Fig. 5f). Nevertheless, 83.6% of the cells in the DN4 unusual cluster express *Trbc1/2* transcripts and of these 70.7% express *Lck* and/or *Ptcra* confirming the T lineage origin of a large fraction of the cells (Supplemental Information File 2). Moreover, there is maintained expression of progenitor drivers (*Kit*, *Lyl1*, *Ezh1* and *Id2*) as well as *Spi1* coding for PU.1 that operates at the critical decision checkpoint determining myeloid or T cell lineage specification. These observations are consistent with not having passed through the preTCR checkpoint as is the maintained expression of early lineage and γδ T cell-linked developmental transcripts such as *Fcer1g*, *Icos*, *Il18rap*, etc. (Extended Data Fig. 3a). The high representation of these unusual cells is not part of the normal ILC or γ/δ T cell development, else they would also appear in the MHC^+^ DN4 library that harbours a similar ILC-γ/δ T cell cluster. Furthermore, the MHC^-^ unusual cells are in a cycling state with high histone transcript expression, together with high expression of *AY036118* (Fig. 3a, 3b), a lncRNA (XR_877120.4) on Chr17 implicated in regulation of thymocyte proliferation possibly mediated by telomeric association^28, 29^. The volcano plot identifies transcripts of high fold-change and probability averaged across the whole cluster hence significance can be driven by a well-represented subset of cells rather than the complete cluster population. Examining select transcripts that are regulated in the same direction in most cells within a cluster, in addition to *AY036118*, the histones represented here by *Hist1h1d* as well as *Lars2* coding for mitochondrial leucyl-tRNA synthetase 2, a marker of high metabolic activity stand out^30^ (Fig. 3b).

Unexpectedly, this analysis led to identification of irregularities in *Rag1* and *Rag2* transcription in the DN4 unusual population where both Rag transcripts are minimal (Fig. 3b). *Rag1* is well expressed in MHC^+^ and MHC^-^ DN3b/4 clusters. *Rag2* is expressed well in the MHC^+^ DN3b/4 cluster while expression in the MHC^-^ DN3b/4 is comparable to that of the DN4 unusual cluster. These findings not only illuminate possible differential regulation of the *Rag1* and *Rag2* transcripts but also show that the MHC^-^ DN3b/4 cells are already experiencing transcriptional aberrations despite appearing phenotypically identical with MHC^+^ DN3b/4 cells. Reduction of Rag1/Rag2 heterodimeric protein activity in the DN4 unusual population due to regulation of *Rag2* transcripts might contribute to the loss of diversity in the β chain repertoire at this stage (Fig. 2d).

## Dysregulated transcriptome of MHC^-^ DN4

Further refinement of the properties of the MHC^-^ DN4 unusual cluster are revealed by single cell β clonotype analysis. Examination of the MHC^+^ DN4 library for the top 20 clonotypes based on cellular representation (Extended Data Table 3a) shows that the majority localise to the DN3b/4 cluster as expected, given appropriate preTCR signalling with minimal tracking to other clusters (Fig. 3c). Similar analysis of the MHC^-^ DN4 library exposes a starkly different distribution where the majority of highly represented β clonotypes map to the DN4 unusual cluster (Fig. 3d). Of the 17 clonotypes represented in the DN4 unusual cluster, for 14 we can identify related cells bearing the same clonotype in the DN3a/3b and DN3b/4 clusters (Fig. 3d right panel). Consequently, we propose that in the absence of pMHC, some cells may differentiate from DN3a through to DN4 but deviate from the normal transcriptome trajectory to map to the unusual cluster. Cell representation of the top 20 clonotypes in the DN4 libraries, normalizing for differences in initial library size, shows 3.25 ± 0.55 cells for each MHC^+^ clonotype (only 2/20 found in the unusual cluster) compared with 6.56 ± 2.07 cells for each MHC^-^ clonotype (17/20 in the DN4 unusual cluster, P < 0.0001). This confirms the increased proliferation implied by the transcriptome signature of the MHC^-^ developing cells in this unusual cluster.

Eight of the top 20 clonotypes are found in the γ/δ T/ILC cluster and five of these are shared with the MHC^-^ DN4 unusual cluster implying T lineage developmental options may remain open without delivery of appropriate preTCR-pMHC-dependent regulatory signals. Of interest is the observation that cells expressing the same unique β clonotype, particularly those MHC^-^ developing cells, tend to group closely together within the UMAP cluster implying conservation of the transcriptional signature, even for occasional clonotypes split between clusters (Extended Data Fig. 6).

The transcript signature of the DN4 unusual population, with upregulation of early progenitor proliferative genes and of *Spi1* controlling the myeloid/T lineage decision point at the DN2a/DN2b transition, connotes a de-differentiation of the DN4 cells in the absence of appropriate preTCR signalling. To examine the possibility that this uncommon transition may generate a transcriptional landscape consistent with aberrant transformation potential, we performed single sample gene set expression analysis (ssGSEA) against cancer modules followed by more refined comparisons with clinically defined T-ALL gene sets. By ssGSEA analysis, the DN4 “unusual” cluster shows a strong score (>1000) against 9 of the top 10 modules defined by maximal score difference from the DN3b/4 clusters of both the MHC^+^ and MHC^-^ DN4 libraries (Fig. 3e). Leukaemia/lymphoma transcriptomes show significant co-ordinated regulation with all 9 of these gene set modules. In contrast, the DN3b/4 clusters of both MHC^+^ and MHC^-^ DN4 libraries tracked together and showed weaker association or even inverse correlation. Further refinement of this analysis compared expression in the DN4 unusual population with published transcript panels defining T-cell acute lymphoblastic leukaemia (T-ALL) focussing upon Early T-cell Precursor ALL (ETP-ALL), a subset of T-ALL with poor prognosis in humans and believed to develop from early thymic progenitors immigrating from the bone marrow^31, 32^. The selected transcripts were grouped as being common to T-ALL generally, or alternatively, representing DN1/2a ETP-ALL prior to committing to the T lineage (“early”), or DN2b ETP-ALL after commitment to the T lineage (“late”). Transcript representation within the MHC^-^ DN4 unusual cluster subsequently was compared with that in MHC^+^ DN3b/4 cells following the expected developmental trajectory (Fig. 3f). Seven of 10 transcripts representing the common panel trended to upregulation, while three showed no change. Except for *Spib* that is weakly upregulated, none of the transcripts in the DN2b-ALL “late” panel were upregulated. In contrast, 5 of the 8 selected genes in the DN1/2a-ALL “early” panel were significantly upregulated, and the remainder all trended upwards, compatible with the cells dedifferentiating from a DN4 state back towards the early progenitor state. As no significant differences were noted in CDR3 length or hydropathy between DN4 Vβ clonotypes developing on MHC^+^ versus MHC^-^ stroma (^8^ and Supplemental Information File 3), the abnormal transcriptome likely emanates from lack of preTCR ligation by MHC and not aberrant preTCR sequences *per se*.

## Abnormal DP subset with dedifferentiation

The DN4 unusual cluster forms one section of a bipartite UMAP cluster that also includes a unique population found only in the MHC^-^ DP thymocyte-like library leading to its classification here as abnormal (Fig. 1b, c). This DP population projects away from the DN4 component due to the expression of *Cd4*, *Cd8a* and *Cd8b1* (Extended Data Fig. 5g) but maps to the same DN4 cluster projection due to the strong expression of *AY036118*, histones, early progenitor-related transcripts, *Spi1* driving non-T lineage commitment in the early DN stages and markers not strongly expressed in the MHC^+^ developing DPbl or DPsm clusters (Fig. 4a). Remarkably, the most significantly upregulated transcripts in this DP abnormal population are transcripts that define the myeloid lineage: *Mpo* (myeloperoxidase), *Prtn3* (proteinase 3), *Ctsg* (cathepsin G), and *Elane* (neutrophil elastase), where expression is specific to this cluster without expression in any of the DN3a to DPsm clusters representing the expected developmental trajectory or in the MHC^-^ DN4 unusual cluster (Extended Data Fig. 5h). Selecting transcripts that are upregulated throughout the DP abnormal cluster confirms the signature *AY036118* profile, cell cycling and DNA packaging using *Hist1h1d* as representative of a broad spectrum of histones, *Plac8* as an oncogenic driver as well as the key myeloid markers, *Prtn3* and *Mpo* (Fig. 4b). *SPI1*, *LYL1*, *LMO2* and *MEFC2* are dominant components of a panel defining human ETP-ALL^33^ and the mouse homologues are upregulated in the DP abnormal cluster, where *Lyl1* and *Spi1* are upregulated above that already seen in the DN4 unusual cluster (Fig. 3a, 4a). Supporting origin from αβTCR T cell lineage, the *Spi1*^+^ cells in the MHC^-^ DP cluster express β variable region transcripts (Fig. 4g) with fully recombined clonotypic TCR β chains in more than 37% of those cells (Extended Data Fig. 5i).

**Fig. 4.**
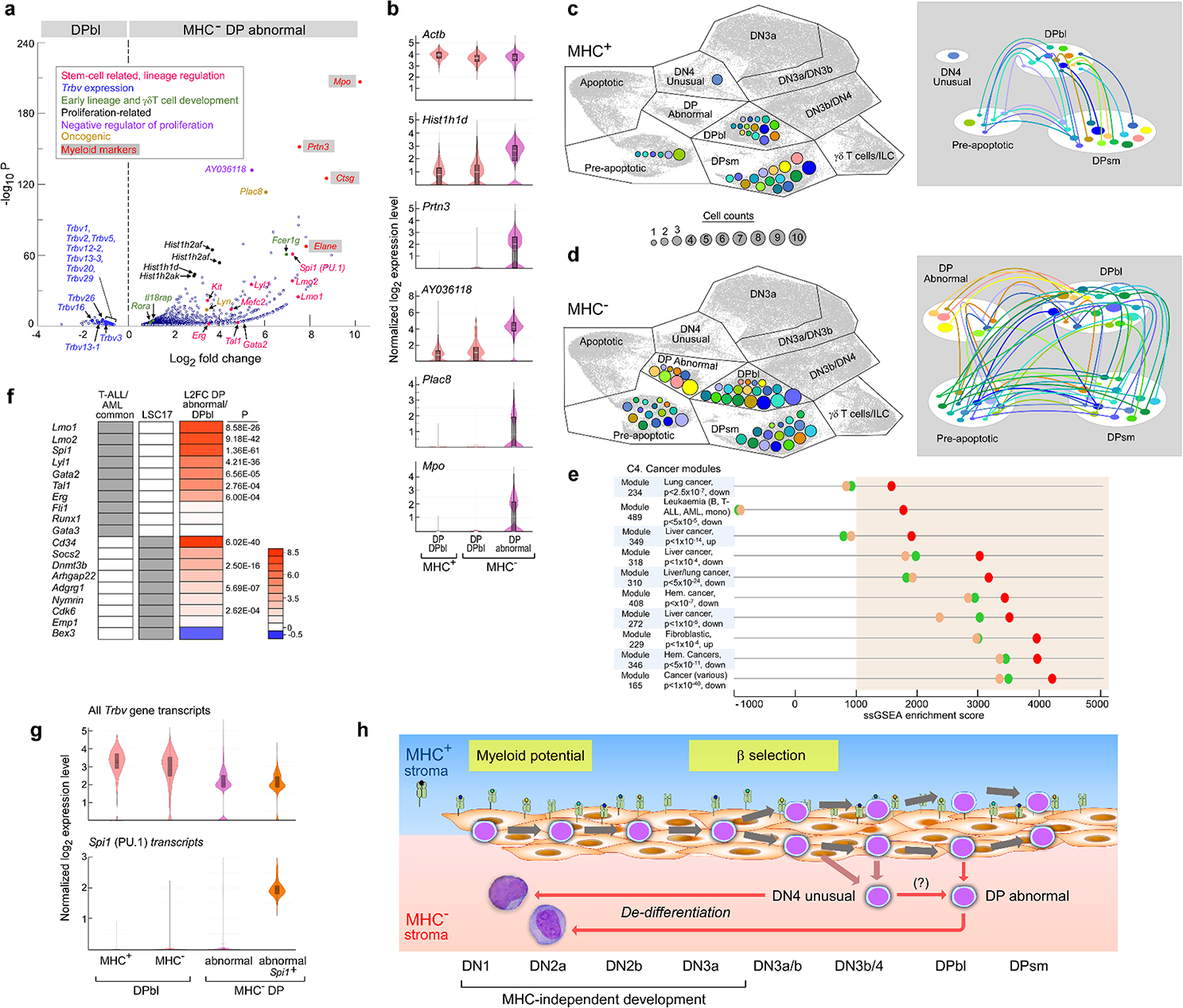
Single cell transcriptomics of the MHC^-^ DP abnormal cluster disclose dedifferentiation and reprogramming to include a myeloid programme. **a.** Volcano plot of significant transcript differences (P<0.05) between the DP abnormal population and the DPbl cluster in the same MHC^-^ DP library. DP abnormal upregulated transcripts shown to the right of zero, downregulated to the left. Functional significance of the highlighted transcripts is listed in the inset box. An additional 10 *Trbv* transcripts were downregulated but did not meet the P<0.05 threshold. **b.** Violin plots depicting indicated log-normalised transcript levels selected for representation within most cells of the DP abnormal cluster that are significantly different to representation with the DPbl clusters within the MHC^+^ and MHC^-^ libraries, respectively. *Actb* transcript is included as a reference. Cluster colours consistent with those depicted in Figs. 1 and 2. P values as in panel a. **c.** Cluster distribution of top 20 DP clonotypes developing on MHC^+^ stroma (left panel). Each colour represents a unique clonotype and circle diameter is proportional to the cell number expressing that specific β chain. Colour/ clonotype specification is unique to the panel and bears no relation to colour use in Fig. 4d or Fig. 3c, d. Right panel depicts clonotype tracking from cluster to cluster maintaining colour concordance with left panel. **d.** DP cluster distribution of top 20 clonotypes developing on MHC^-^ stroma. Each colour represents a unique clonotype and circle diameter is proportional to the cell number expressing that specific β chain (see cell count scale common to Fig. 4c and Fig. 4d). Right panel depicts clonotype tracking from cluster to cluster. **e.** ssGSEA scores for MHC^+^ DPbl (green), MHC^-^ DPbl (light orange), and MHC^-^ DP abnormal (red) cells analysing module gene sets with co-ordinated gene regulation in tumours (MSigDb C4) and probability of association of specific tumors with up- or down-regulation of the specific module. Significance of ssGSEA scores as described in Fig. 3e legend. **f.** Comparison of MHC^-^ DP abnormal cluster with the DP cluster developing on MHC^+^ stroma for transcripts reported as dysregulated in human AML (T-ALL/AML common) or as overexpressed in CD34^+^ leukaemic stem cells (LSC) from AML patients (LSC17). Note that not all 17 transcripts in the human LSC17 panel have equivalents in the mouse while others are expressed at levels too low (*i.e.* 0) by scRNA-Seq to generate a meaningful fold-change ratio. Column 3 heatmap depicts log2-fold difference for the indicated transcript (scale to right); P indicates significance. If no value indicated, then P>0.05). **g.** Co-expression of *Spi1* and *Trbv* transcripts in *Spi1*^+^ MHC^-^ DP abnormal cells. **h.** Schematic of proposed dedifferentiation for thymocytes developing in the MHC^-^ condition (pink area). Proliferation and development appear identical with that in the control MHC^+^ condition (light blue area) as cells progress from early progenitors into the DN3a compartment. The MHC presents a diverse range of small self-peptides. Following absence of preTCR binding to pMHC ligand, however, multiple abnormalities in β chain representation develop as noted in the text, apparent within DN4 unusual cells found with minimal representation in the MHC^+^ libraries (reddish-brown arrows) as well as in a new population, DP abnormal, absent in the MHC^+^ DP library.

To investigate the possibility that the DP abnormal cluster arose from aberrant expansion of one HSC in the progenitor pool, the fraction of Chr:Y^+^ cells in each cluster was determined. These results do not support stochastic growth independent of stromal cell MHC expression (Supplemental Information File 6). Instead, comparison with a matched panel of autosomal genes reveals that the transcriptional abnormalities are found equivalently in Chr:Y^+^ and Chr:Y^-^ cells within the MHC-DP abnormal cluster. Further strengthening the proposal that the DP abnormal cells are following a path deviating from the wild-type developmental pathway, 80% of the *Mpo^+^* cells and 84% of the *Mpo^+^Spi1^+^* cells are co-expressing *Lck* and/or *Cd3e* (Extended Data Fig. 5j).

Both NKT cells and Mucosal-associated invariant T cells (MAIT) develop from the DP population but there is no evidence that these cells are developing as an alternative path to canonical αβ TCR cells in the absence of pMHC ligation based on two orthogonal findings in our data. First, their respective transcriptional signatures do not map to the DP abnormal population (*Rorc*, *Tbx21* and *Gata3* for NKT; *Zbtb16*, *Drosha*, and *Il18* for MAIT^32, 34^, although *Mr1* is 2-fold upregulated). Second, all 3 TCR β chains restricted to mouse NKT cells (*Trbv1*, *Trbv13* alleles and *Trbv29*) are downregulated in the DP abnormal population while those β chains restricted to mouse MAIT cells are downregulated (*Trbv13* alleles) or unchanged (*Trbv19*) (Fig. 4a)^34^.

As observed for the DN4 libraries, the distribution of the top 20 clonotypes by cell representation was markedly different between the MHC^+^ DP library and the MHC^-^ DP library (Extended Data Table 3b). Average cell representation of each clonotype was higher in the MHC^-^ condition (Fig. 4c, d), and tracking showed that this difference was retained in the DPsm and pre-apoptotic clusters. Representatives of both the DP abnormal and of the DPsm are found in the DPbl population, but there is minimal overlap between the DPsm and DP abnormal cells implying cluster destiny is specified at the DPbl stage. Given the proliferative transcript signature, the early progenitor profile, and the presence of the myeloid markers in the context of a DPbl population origin, we examined the transcriptome by ssGSEA for evidence that the DP abnormal population may be entering into a state conducive to future myeloid dysplasia or leukaemia development (Fig. 4e). MSigDb C4 cancer module 489 generated the highest differential score, a cell profile that is strongly associated with leukaemias including T-ALL and AML. The signature panel for regulatory gene abnormalities in CD34^+^ leukaemic stem cells isolated from acute myeloid leukaemia (AML) patients overlaps completely with that for T-ALL (Fig. 3f, 4f)^35^. A further AML CD34^+^ leukaemic stem cell panel (LSC17)^36^ was used to assess any potential equivalence of the DP abnormal cells with transformed AML leukaemic stem cells (Fig. 4f). The DP abnormal cells expressed 7/10 signature transcripts in the T-ALL/AML common panel at levels significantly higher than developing DPbl cells. The minimal change in *Gata3* and *Runx1* may indicate the ongoing T cell lineage programme in both subpopulations. Comparison with representatives of the human LSC17 panel found significant upregulation of 4/9 markers with a further 3 trending upward. *Cd34*, the canonical haematopoietic stem cell marker, was the most profoundly upregulated (293-fold). The high expression of *Cd34* coupled with persistence of *Erg* (Fig. 1a) in the MHC^-^ DP abnormal population, absent in the MHC^+^ libraries, points to an earlier progenitor environment in the absence of pMHC-driven preTCR signalling.

## *B2m* and *H2-Ab1* dKO preTCR signalling

We next examined gene expression in the thymus of MHC^+^ B6 mice and mice on the same background carrying double knockout (dKO) mutations for both *B2m* and for *H2-Ab1* previously created to abrogate expression of MHCI and MHCII^21^ (MHC^-^). We ascertained whether the phenomena observed *in vitro* were recapitulated *in vivo*. Cell recoveries indicated a significant increase in DN3a cells in the MHC^-^ thymi, a differential that extended less significantly through the DN3b to DN4 stages (Extended Data Fig. 7a). Examining gene expression for cells transitioning from the DN3a to immature CD8 single positive (ISP) stage (Supplemental Information File 4), strong downregulation of *B2m* and moderate downregulation of *H2-Ab1* was observed for H-2 negative thymocytes (Extended Data Fig. 7b). For the transcript changes occurring in the DN3a to ISP transition depicted in Fig. 1a, however, we found no difference between MHC^+^ and the MHC^-^ thymocytes (Extended Data Fig. 7c). Moreover, no reduction in the *Trbv* transcription at the DN4 stage was observed in the MHC^-^ thymocytes (Extended Data Fig. 7d). In contrast, a defined hallmark of preTCR signalling, the upregulation of anti-apoptotic *Bcl2a1* family transcripts, was clearly observed in the MHC^+^ but not detected in MHC^-^ DN4 thymocytes (Extended Data Fig. 7e), while canonical *Bcl2* pathway transcripts were similar in both^37^. Upregulation of *Trav* transcripts dependent upon preTCR signalling was significantly stronger in the MHC^+^ than in MHC^-^ mice (P=2ˣ10^-8^; Extended Data Fig. 7f)^38^. Of interest, the *Pim1* proto-oncogene associated with foetal haematopoiesis and overexpressed in myeloid and lymphoid leukaemias^39^ is one of the strongest expressed transcripts detected in the MHC^-^ libraries but barely detected in the MHC^+^ libraries (Extended Data Fig. 7e). Analysis of complete β chain repertoires for the entire thymus representation of DN3a to ISP cells was uninformative; for each library more than 98.9% of the clonotypes were represented by 3 or fewer UMI leading to such high repertoire diversity scores that no significant differences were observed between libraries.

## Compensatory MHCIb upregulation in dKO mice

Remarkably, in all MHC^-^ libraries, both *H2-T3* (TL) and *H2-T22* were dramatically upregulated over those in MHC^+^ libraries (Extended Data Fig. 7b; *H2-T3* mean Fragments Per Kilobase of transcript per Million mapped reads (FPKM) for MHC^+^ = 0.75 and for MHC^-^ = 44.1 FPKM; *H2-T22* mean FPKM for MHC^+^ = 21.0 FPKM and for MHC^-^ = 97.0). In contrast, OP9-DL4 *H2-T22* expression was similar between the MHC^+^ and MHC^-^ cells and *H2-T3* (TL) was undetectable in either of the isogenic stroma (Extended Data table 2). Also, noteworthy in MHC^-^ libraries was upregulation of *H2-Q10* and *H2-T-ps*, the latter now believed to be protein coding (NCBI Gene ID: 667803).

The enhanced transcription of *H2-T22*, *H2-T3* (TL), *H2-Q10* and *H2-T-ps* genes implies that adaptation *in vivo* maintains functional β selection by upregulating non-classical minor MHCIb products, thereby compensating for loss of classical MHCIa alleles. This phenomenon is not operative in the OP9 cultures. Our animal studies are not only consistent with the apparent normal phenotypic thymocyte development of *B2m* and for *H2-Ab1* double knock out mice observed previously^21^ but underscore the complexity of vital *in vivo* biological signalling including mechanisms to override pathway blockade via compensatory adaptation. Nonetheless, preTCR signalling is not entirely normal as evidenced by the failure to observe upregulation of *Trav* and *Bcl2a1,* in agreement with the suggestion that the narrow width of the MHC1b α1α2 presenting platform relative to that of MHC1b might attenuate preTCR signaling^8^. Given this biological readjustment *in vivo,* the utility of synchronous *in vitro* culture to pinpoint critical developmental steps is essential.

## preTCR-pMHC safeguards orderly development

The current *in vitro* study reveals that preTCR-pMHC interactions sculpt the transcriptome of DN3 and later stage thymocytes, in addition to fostering β clonotype diversity in the αβ T cell lineage. By comparing single cell transcriptomes of a pool of B6 DN3 foetal thymocyte progenitors differentiating in parallel on isogenic MHC^+^ and MHC^-^ OP9-DL4 epithelial stroma, three irregular UMAP clusters were uncovered on the MHC^-^ stroma. The *first*, an aberrant DN3b-DN4 transitional population, lacked evidence of preTCR signalling but maintained Notch signalling and manifest a broad decrease in *Trbv* transcripts. The *second*, a DN4 unusual population, minimally present in the MHC^+^ population, abnormally upregulated genes involved in earlier stages of thymic renewal (*Lyl1*, *Kit*, *Id2*, *Dtx1* and *Bcl11a*), T cell co-stimulatory function (*Icos*), adhesion function (*Itgb3*) and cytokine receptors involved in inflammation (*Il18r*, *Il23r*). The *third,* an entirely anomalous cluster, DPbl abnormal, expressed *Cd4*, *Cd8a* and *Cd8b1*, and simultaneously multiple myeloid genes (*Mpo*, *Prtn3*, *Ctsg*, *Elane*, *Hdc* and *Cst7*). Both DN4 unusual highly proliferating cells and DPbl abnormal myeloid-like cells expressed the *AY036118* gene implicated in control of thymocyte proliferation^29^. Thus, even in short-term *in vitro* culture of progenitors, striking deviations in normal progeny arise in the absence of preTCR-pMHC ligation.

Such aberrations of developmental programs at DN and DP thymocyte stages are noteworthy given that human T-ALL represent malignancies of these same phenotypic subpopulations including a subset of DN early T cell precursors^31, 40^. Collectively, this aggressive group of malignancies results from key genetic abnormalities including instabilities fostering rearrangements and/or deletions of *TCRB*, *TCRA* and *TCRD* loci, genes linked to cell cycle growth control (*Cdkn2a* or *Cdkn2b*) and mutations associated with hyperactive Notch signalling^41, 42^. The latter are present in ≥50% of cases, often with additional mutations of transcription factors and signalling pathways^43^.

*In vivo* mouse over-expression of transcription factors (*Tal1* and *Tlx1*) also results in T-ALL^44, 45^ with acceleration of disease mediated by additional mutations such as those involving *Bcl11b* or *Notch1*^46, 47^. Over-expression of Notch intracellular domain, the signalling entity generated by normal ligation of Notch and proteolytic cleavage by γ-secretase, in early extraembryonic yolk sack haematopoietic precursors transplanted into lethally irradiated mice led to T-ALL with a DP thymocyte blast phenotype^48^. Perhaps even more strikingly, disruption of competition between “new” bone marrow-derived immigrants and “existing” DN3a thymic resident progenitors leads to aberrant self-renewal of the latter culminating in murine T-ALL reminiscent of human T-ALL in virtually all respects, replete with their development of activating *Notch1* mutations^49^. These DN3a thymic self-renewal progenitors give rise to TCRβ-deficient DP thymocytes with a high frequency of non-productive β gene rearrangements expressing *Notch1* and *Ptcra* transcripts consistent with ongoing Notch signalling^50^.

The DP blast abnormal cluster’s expression of myeloid genes and transcriptional signatures shared with AML and ETP-ALL stem cells^35, 36^ suggest that a subset of myeloid malignancies may arise from the DP compartment after further transformation, particularly in light of the clinical entity of mixed phenotype acute leukaemia (MPAL) expressing both lymphoid and myeloid malignant markers simultaneously^33, 51^. Thus, consideration needs to be given to the possibility that rather than singularly arising from ETP, a thymic genesis of certain haematopoietic malignancies can involve de-differentiation from later stages of development including DP thymocytes and maybe even involve transdifferentiation to other lineages. De-differentiation is a normal process whereby cells progress in a retrograde manner from a more to a lesser differentiated state as a safeguard against progenitor loss^52^. Although such phenomena have been induced by chemical or genetic means in the haematopoietic system^53–, 55^, here we demonstrate that lack of appropriate signalling during development leads to reprogramming. The UMAP projection localizes the abnormal cluster cells between the expected developmental path and the apoptotic cluster. While detected within a synchronised window of differentiation *in vitro*, rapid *in vivo* removal of apoptotic cells by phagocytes in the thymus would obscure the destiny of the unusual and abnormal cluster cells.

We postulate that self-pMHC reactivity triggers preTCRs on thymocytes during β-selection, attendant downregulation of Notch signalling, modulation of cell-cell adhesion, migration, and metabolism. Recent immunofluorescence microscopy studies demonstrated formation of an immunological synapse between DN3a thymocytes and stroma, thereby creating a preTCR platform around β-selection to integrate cues involving Notch ligand, the CXCR4 ligand as well as pMHC on thymic stroma likely involving asymmetric cell division to foster further differentiation^56, 57^. A cellular niche of this type could serve as a pivotal nexus within the developmental circuit to terminate cellular plasticity and foster orderly downstream development. The preTCR on DN3 thymocytes and pMHC on stroma may offer bidirectional signalling in receptor and ligand expressing cells, given precedent in other systems^58^. This circuit can go awry, however, if preTCR-pMHC ligation falters because of absent functional ligands, should there be disruption of the physiologic regulation of attendant associated pathways, or if there is dysregulated entrance of progenitors into and/or exit from their developmental niche. In this view, the previously unexpected oncogenic potential of a preTCR lacking its Vβ variable domain to induce DP T cell lymphoma now can be rationalized given that the Vβ domain is the only receptor domain capable of binding to pMHC^59^. Likewise, generation of intra-thymic AML (^60^ and references therein) can be understood as a possible consequence of early developmental plasticity and thymic niche anomalies. TCR gene rearrangement processes necessary for T-lineage repertoire formation bracket the β-selection that fosters clonal expansion and repertoire diversification, thereby creating a further vulnerability for tumorigenesis. Somatic TCR repertoire formation affording protective adaptive immunity incurs this potential cost. Lastly, our findings emphasize that while thymocyte progression *per se* can occur in the absence of classical MHC ligand-dependent preTCR function, those self-pMHC interactions are essential for normal development and to mitigate aberrant de-differentiation. The upregulation *in vivo* of non-classical MHCIb in the double knockout mice to preserve ligand-dependent preTCR function underscores this biology.

## Methods

### Mice

Six-week-old C57Bl/6 (B6) and B6.129-*H2-Ab1^tm1Gru^ B2m^tm1Jae^* N17^21^ (MHC^-^) mice were purchased from Taconic Farms Inc. and housed at the DFCI Animal Facility, accredited by the Association for Assessment and Accreditation of Laboratory Animal Care (AAALAC). All maintenance, breeding, and experimental procedures were approved under Dana-Farber Cancer Institute Institutional Animal Care and Use Committee (IACUC) protocols 03-138 and 04-113. Euthanasia was by CO_2_ inhalation followed by cervical dislocation. Following removal from the uterus, E14.5 fetuses were euthanized by decapitation with surgical scissors. Where appropriate, no gender preference was expressed for experimental animal use.

### Reagents

The OP9-DL4 parental (MHC^+^) cell line, and the MHC^-^ and scH-2K^b^ variants, were developed and used as described previously^7, 60^. Anti-mouse CD44-APC/Cy7 (clone IM7) and anti-mouse CD117-APC (c-Kit; clone 2B8) were obtained from BD Biosciences. Anti-mouse CD24 and anti-mouse CD24-FITC (clone M1/69), anti-mouse CD3e-BV605 (clone 145-2C11), anti-mouse CD4-Pacific Blue and CD4-BV711 (clone RM4-5), anti-mouse CD8a-PerCP/Cy5.5 (clone 53-6.7), anti-mouse CD8b.2-PE (clone 53-5.8), anti-mouse CD11b-biotin (clone M1/70), anti-mouse CD11c-biotin (clone N418), anti-mouse CD19-biotin (clone 6D5), anti-mouse CD28-PE (clone E18), anti-mouse CD45-APC (clone 30-F11), anti-mouse NK1.1-biotin (clone PK136), anti-mouse Gr-1-biotin (clone RB6-8C5), anti-mouse Ter-119-biotin (clone TER-119), anti-mouse TCRγ/δ (clone GL3), streptavidin-BV421, an Zombie Aqua were obtained from Biolegend. Anti-mouse Ly-6A/E (Sca1)-FITC (clone D7) and anti-mouse CD25-PE/Cy7 (clone PC61.5) were obtained from eBioscience.

### Analysis of B6 thymocyte-like development *in vitro*

Isolation of wild-type haematopoietic stem cells (HSC) followed the procedure described previously^7^. Briefly, fetal liver cells from 30 E14.5 B6 embryos from 3 dams were depleted of B cells using anti-CD24 and complement lysis (Cedarlane) followed by staining with anti-CD4-Pacific Blue, anti-CD8-PE, anti-ScaI-FITC and anti-CD117(c-Kit)-APC. The CD4^-^CD8^-^ (lineage negative, lin^-^) ScaI^+^ c-Kit^+^ cells (HSC) were isolated by a Becton-Dickinson FACS Aria II cell sorter. For the T cell repertoire analysis from pooled FACS-sorted thymocyte-like cells, 2,000 HSC were seeded onto 70 to 90% confluent layers of wild-type OP9-DL4 cells (MHC^+^), or onto MHC^-^negative OP9-DL4 cells (MHC^-^), or onto the MHC^-^ cells re-expressing a single chain H-2K^b^ presenting VSV8 peptide (scH-2K^b^), in six-well plates (i.e. six independent cultures) in α-MEM without nucleosides + 15% FCS (OP9 media), HEPES (10mM), and gentamycin supplemented with Flt3 (5 ng/ml; R&D) and IL-7 (1 ng/ml; Peprotech). For the scRNA-seq experiments, 30,000 similarly prepared HSC were seeded onto MHC^+^ or MHC^-^ stromal cells under the same conditions, increasing the replicates to ten 10 cm dishes/OP9 variant. After growth for 9 days, cells were isolated from the cultures and counted prior to FACS separation to enrich by surface antigen phenotype for cells at different stages of thymocyte-like differentiation.

### Cell sorting, library preparation and data processing for scRNA-seq and TCR V(D)J repertoire characterization

For scRNA-seq analysis, cells were stained with a cocktail consisting of Zombie Aqua for gating of non-viable cells, anti-CD45-APC for gating of haematopoietic cells, with biotinylated anti-CD11b, anti-CD11c, anti-NK1.1, anti-mouse TCRγ/δ, anti-Gr-1, anti-Ter119, and anti-CD19 followed by streptavidin-BV421 for gating of non-T lineage cells, and of anti-CD4-BV711, anti-CD8α-PerCP/Cy5.5, anti-CD44-APC/Cy7, anti-CD25-PE/Cy7 and anti-CD28-PE for gating and collection of DN3a, DN3b, DN4 and DP thymocyte-like cells on a FACS Aria II cell sorter (Extended Data Fig. 1). Note that residual ILC-γ/δ-like T cells in the DN4 subset represent cells with a ILC precursor (*Id2*, *Zbtb16*), ILC2 (*Gata3*, *Rora*), γ/δ T cell-like transcriptome but with no or low surface TCR expression. For each condition (MHC^+^ or MHC^-^), 50,000 DN3a, DN3b, DN4 and DP cells were collected by FACS for application to a 10X Chromium controller (10X Genomics) and recovery of 8,932 ± 920 (mean ± s.e.m.; n = 8) processed cells for gene expression (5’ GEX) and TCR V(D)J sequence library construction. Bar coding and 5’ library construction using v1.0 chemistry was performed precisely following the manufacturer’s protocol. Targeted mouse TCR recovery utilised the Chromium Single Cell V(D)J Enrichment Kit for mouse T cells. All libraries were single i7-indexed using the Chromium i7 Multiplex kit.

Following isolation and clean-up of library DNA, integrity was assessed using an Agilent Bioanalyzer and quantification by Qubit analysis (Invitrogen). All libraries were adjusted to ∼50 ng/µL, where peak fragment size (including Illumina adapters) for the gene expression (5’ GEX) libraries averaged 473 bp and ranged from 300 – 740 bp for the 5’ TCR libraries representing ongoing recombination products in the developing thymocyte libraries. Sequencing (150 PE) was performed on HiSeq 3000 utilizing 4 lanes where two 5’ GEX libraries (2 x 40% of reads) and two TCR libraries (2 x 10% of reads) were sequenced per lane. Following conversion of the *bcl2* sequencing files to *fastq* format, the 5’ GEX sequencing results were pipelined to Cellranger 3.1.0 using the GRCm38.p6/mm10 mouse genome as reference and the TCR files were pipelined to Cellranger V(D)J 3.1.0 using vdj_GRCm38_alts_ensembl-3.1.0.gz-3.1.0 as reference, all using default parameters. Gene expression data from all libraries were aggregated by Cellranger to generate a UMAP of all libraries into the same 2D space. For aggregation, the count output files for each Chromium controller well were processed using the “aggr” command to produce a single feature-barcode matrix containing all the data. Since barcodes may overlap between libraries, a well suffix is added to each barcode-nucleotide sequence to hardcode well origin. Before merging, depth normalization is performed to subsample reads for each library to equalize the number of reads confidently mapped to the transcriptome. Prior to Principal Component Analysis, the UMI counts were normalized towards the median across all cells by multiplying each cell’s UMI count by a scaling factor of the median UMI count across all the cells divided by the UMI count for the cell. The matrix is log-transformed then centered and scaled per-gene such that the mean is 0 and the standard deviation is 1 prior to clustering. Consequently, all data used for differential expression is log-normalized and a pseudocount of 1 was added to both the numerator and denominator of the mean expression. For a cluster or selected cell subset within a cluster, log2-fold change was either tested against the mean expression for all other cells (global analysis) or against a selected cluster or subset (local analysis) using the Loupe browser 4.2.0 together with the Loupe V(D)J browser 3.0.0 (10X Genomics) for integration of TCR clonotype parameters.

### Bulk population TCR repertoire protocol and data processing for cells developing *in vitro*

For the bulk population β repertoire analyses of thymocyte-like cells developing on the MHC^+^ and MHC^-^ stromata, respectively, cells were stained with anti-CD45-APC, anti-CD4-Pacific Blue, anti-CD8-PE, anti-CD25-PE/Cy7 and anti-CD44-APC/Cy7 for simultaneous collection of DN3, DN4. DPbl and DPsm thymocytes on a FACS Aria II cell sorter (Supplemental Information File 4). Contaminating OP9 cells expressed GFP permitting their exclusion while selection for CD45 expression ensured only hematopoietic cells were used for subset delineation. Cells were gated as CD4^-^CD8^-^ (double negative, DN) and CD4^+^CD8^+^ (double positive) from which 10,000 cells each of DN3 (CD25^+^CD44^-^) cells, DN4 (CD25^-^CD44^-^) cells, DPbl (blast cells, in cell cycle; CD4^+^CD8^+^ high forward scatter) and DPsm (small, more mature cells; CD4^+^CD8^+^ low forward scatter), were collected. For each population, the cells were immediately deposited into TCL lysis buffer (Qiagen) supplemented with 2-mercaptoethanol (1%) on ice, snap-frozen by immersion in dry-ice-methanol and stored at –80°C until processed for RNA extraction and β chain repertoire analysis.

Total RNA was extracted from each sample of 10^4^ cells using the PicoPure column purification system (Applied Biosystems). Subsequently, the procedure followed precisely that described by Mamedov *et al.*^61^. Briefly, using a 3′ *Trbc* (TCRβ constant region) universal primer, 1st strand cDNA was synthesized from the starting RNA and a universal “Switch” primer ligated to the 5′ ends. Nested/extended PCR amplification through the universal ends yielded unbiased amplification of transcripts containing the complete V(D)J region and a 5′ segment of the *Trbc*. In the second PCR, pentanucleotide bar codes were introduced to tag each library with unique barcodes at both 5′ and 3′ ends. Following quality control using the Agilent 2100 Bioanalyzer and Illumina adapter addition, samples were sequenced (150 PE) on the MiSeq platform. Library sequences were deconvoluted from the fastx sequence output files using the barcode splitter module of the FASTX toolkit (http://hannonlab.cshl.edu/fastx_toolkit/index.html). The deconvoluted library sequences were aligned to Vβ regions in the GRCm38.p6/mm10 mouse genome followed by clone assembly and CDR3 extraction using the MiXCR suite running under Java^62^. Output provided V, D, J, and Cβ usage, CDR3 nucleotide and amino acid sequence, sequence quality, and relative representation by read count. The VDJtools analytical package was used to track and compare clonotypes within the libraries^63^.

### Gene expression and total β clonotype analysis of thymus DN3 to ISP cells

The thymus from each of 3 B6 and 3 MHC^-^ mice (all males aged 3 weeks) was isolated and the cells dispersed into RPMI-1640 medium treating each thymus as an individual sample. The cells were incubated with anti-CD4 (clone L3T4) covalently linked to microbeads (Miltenyi Biotec) used at a ratio of 100 µL beads/10^8^ cells then incubated for 10 min on ice. The thymocyte/microbead mixtures were applied to replicate LS MACS columns in a MidiMACS separator and unbound cells collected as CD4-depleted populations removing DP thymocytes and CD4SP thymocytes. The CD4-depleted populations were then sorted to remove the non-T lineage cells as described above (viable, non-T lin^-^ < 0.05% CD4^+^). Following gating on DN cells (CD4^-^CD8^-^), cells were gated further on the DN3/4 population (CD44^-^), and then into three further gates of CD25^hi^CD28^lo/int^ (DN3a), CD25^int^CD28^hi^ (DN3b), and CD25^lo^CD28^hi^ (DN4) as outlined in Supplemental Information File 4. Following gating on the CD8^+^ cells in the viable, non-T lin^-^ population, the cells were further gated on the CD24^hi^CD3^-^ cells (Immature single positive; ISP) and isolated populations collected into TCL lysis buffer as described above. For each mouse this procedure yielded the complete representation of all phenotypically defined DN3a, DN3b, DN4, and ISP thymocytes. Total RNA for each population was prepared using the RNAqueous-4PCR protocol (Applied Biosystems/Life Technologies). From the isolated total RNA, 200 ng was removed for NGS library preparation (SMART-Seq v4 Ultra Low Input RNA, Takara), Illumina adapter addition, and sequencing (PE150, Novaseq platform, ∼40 ͯ 10^6^ reads/sample) for gene expression analysis (Medgenome). The remaining RNA was used for total population repertoire determination following the protocol of Mamedov *et al*^61^. with minor differences to the procedure described above. To reduce errors introduced by PCR amplification as well as estimate individual RNA contributing to a particular clonotype, the “Switch” primer incorporated a region with a universal molecular identifier (UMI) motif of 12 nucleotides within which were interspersed several deoxyuridine nucleotides subsequently treated after cDNA synthesis with uracyl deglycosylase to prevent participation of the Switch primer in the downstream PCR reactions. The individual barcoded DN3a, DN3b, DN4 and ISP libraries for each animal were pooled and Illumina adapters added to generate one total thymus library of these stages for each animal. Following sequencing (PE150, Novaseq platform by Medgenome), library deconvolution, assembly, alignment and UMI processing was handled by the MIGEC package^64^ to determine β clonotype repertoire based on UMI rather than total reads. The output was then pipelined directly to the VDJtools package^63^ as described above. Repertoire diversity was assessed using CalcDiversityStats module of the VDJtools package based on the D50 and Diversity Index (DI)^65^ that yields a value in the 0 – 1 range where 1 = maximal diversity.

### Y chromosome fractional analysis of transcriptionally defined clusters

To address the possibility that the well-represented DN4 unusual population (1776 cells, 14.3% of DN4 library) and DP abnormal population (2512 cells, 19.3% of DP library) in the MHC^-^ condition represent the clonal development of a single or limited number of aberrant progenitor cell(s) during the 9-day culture, skewing of the initial HSC male-to-female cell ratio was assessed by examining the XY cell fraction in each cluster. Given that the initial seeding of 30,000 foetal liver progenitors/initial culture plate originated from a common pool, the ratio of male (XY) cells to female (XX) cells should be maintained across all MHC^+^ and MHC^-^ cultures through to the isolation of phenotypically defined subsets and transcriptionally-defined subsets (clusters) within. Accordingly, a skewed XY/XX ratio within a cluster may indicate non-uniform clonal expansion.

Chromosome Y transcripts likely to be expressed were determined as described in the Supplemental Information File 6 (and ^67, 68^) and the list screened against all libraries to generate a panel of 4 transcripts (*Ddx3y*, *Eif2s3y*, *Kdm5d*, *Uty*) found to be consistently expressed and detectable across all libraries and clusters. The representation of these transcripts was then used to define presence of a Y chromosome. The reverse procedure utilizing transcripts with increased representation in XX cells was not feasible as none of the identified, skewed, transcripts were detected at levels high enough or specifically enough to characterize a cell as definitively XX. Consequently, results were expressed as fraction of cells within a cluster characterized as XY. To exclude observed skewing being a result of apoptotic or other processes occurring in a specific cluster independent of supporting stroma, a panel of genes matched to expression of the Y transcript panel was determined. This latter panel of autosomal gene transcripts (*Cdk8*, *Slc25a5*, *Pank1*, *DFFB*) provided an internal control for cluster-specific skewing within the stroma-specific libraries unrelated to aberrant clonal development (Supplemental Information File 6).

### Statistical analysis

For all the gene expression results, P represents the adjusted p value (P*_adj_*) where P<0.05 was considered significant. Standard parametric statistics followed by Students t test and 2-tailed probabilities were used for all group comparisons. For comparison of small lists of transcripts representing gene expression levels (e.g. *Trbv* alleles), paired t-test or the Chi-square test (utilizing MHC^+^ as “expected”) were used.

## Acknowledgements

This research was supported by NIH NIAID grant AI136301. CMM and PHL were supported additionally by the Expect Miracles Foundation and the Robert and Renée Belfer Foundation. We gratefully acknowledge Dr. Jia-huai Wang for scientific discussion and insight, and Drs. David A. Barbie and Cloud P. Paweletz (Robert and Renée Belfer Center for Applied Cancer Research, Dana-Farber Cancer Institute) for facilitating the scRNA-Seq analyses.

## Author contributions

Conceptualization, J.S.D.-C., A.A., R.J.M., W.H., M.J.L., and E.L.R.; Methodology, J.S.D.-C, A.A., R.J.M., and E.L.R.; Investigation, J.S.D.-C, A.A., C.M.M., and P.H.L.; Writing-Original Draft, J.S.D-C., and E.L.R.; Writing-Review and Editing, J.S.D.-C., R.J.M., A.A., W.H., M.J.L., and E.L.R.; Funding Acquisition, M.J.L., and E.L.R.; Supervision, J.S.D.-C., and E.L.R.

## Data availability

All sequence files deposited in NCBI Gene Expression Omnibus (GEO) under accession GSE186049.

## Additional information

Supplementary Information is available for this paper. Correspondence and requests for materials should be addressed to

## Extended Data

**Extended Data Fig. 1.**
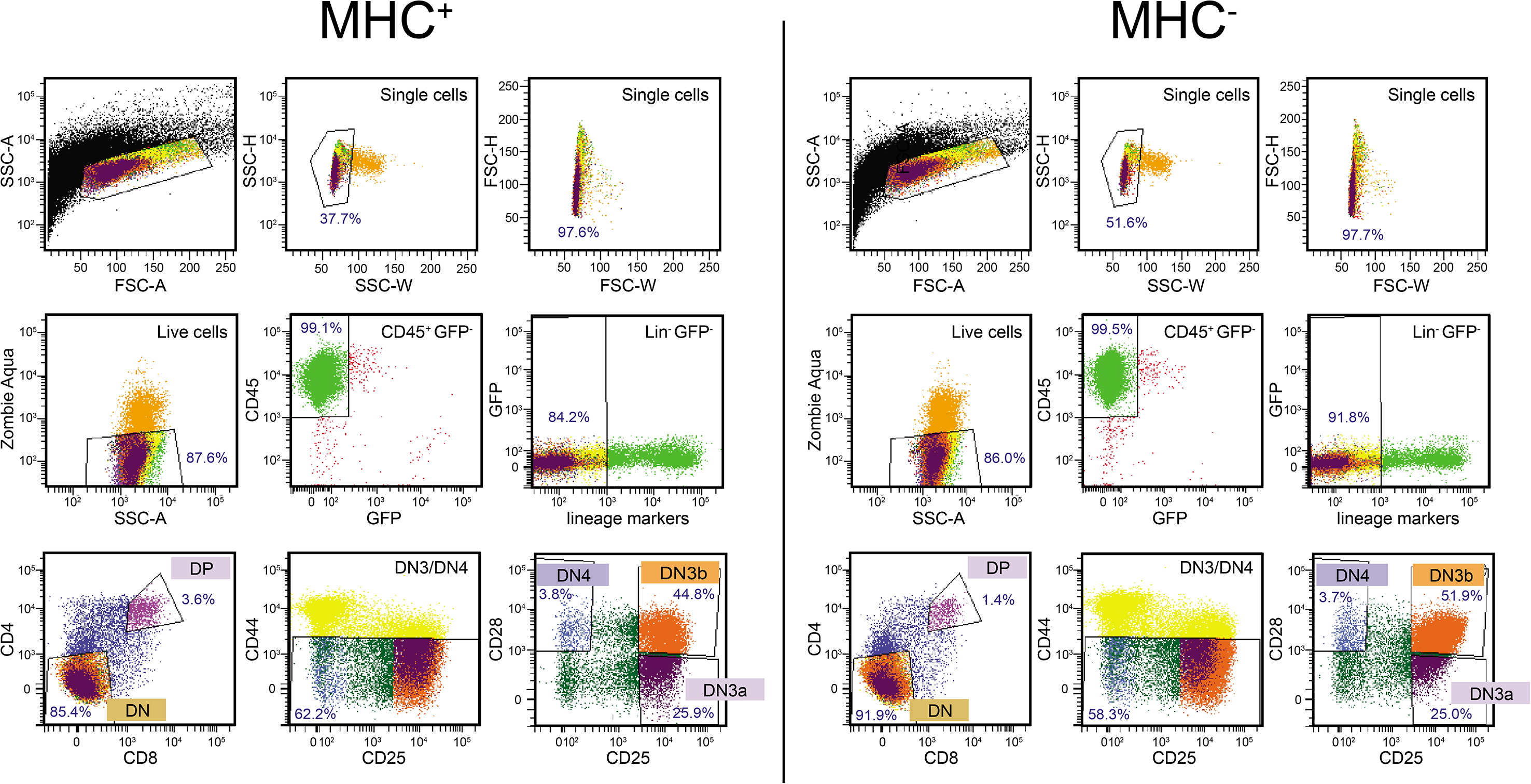
Schematic for FACS isolation of thymocyte subsets (DN3a, DN3b, DN4, DP) for 10X scRNA-Seq and single cell TCR α and β chain clonotype sequencing. Sorted cells were isolated as DN3a cells (CD25^+^CD44^-^CD28^-^), DN3b cells (CD25^+^CD44^-^CD28^+^), DN4 (CD25^-^CD44^-^CD28^+^) cells, and DP (CD4^+^CD8^+^) cells.

**Extended Data Fig. 2.**
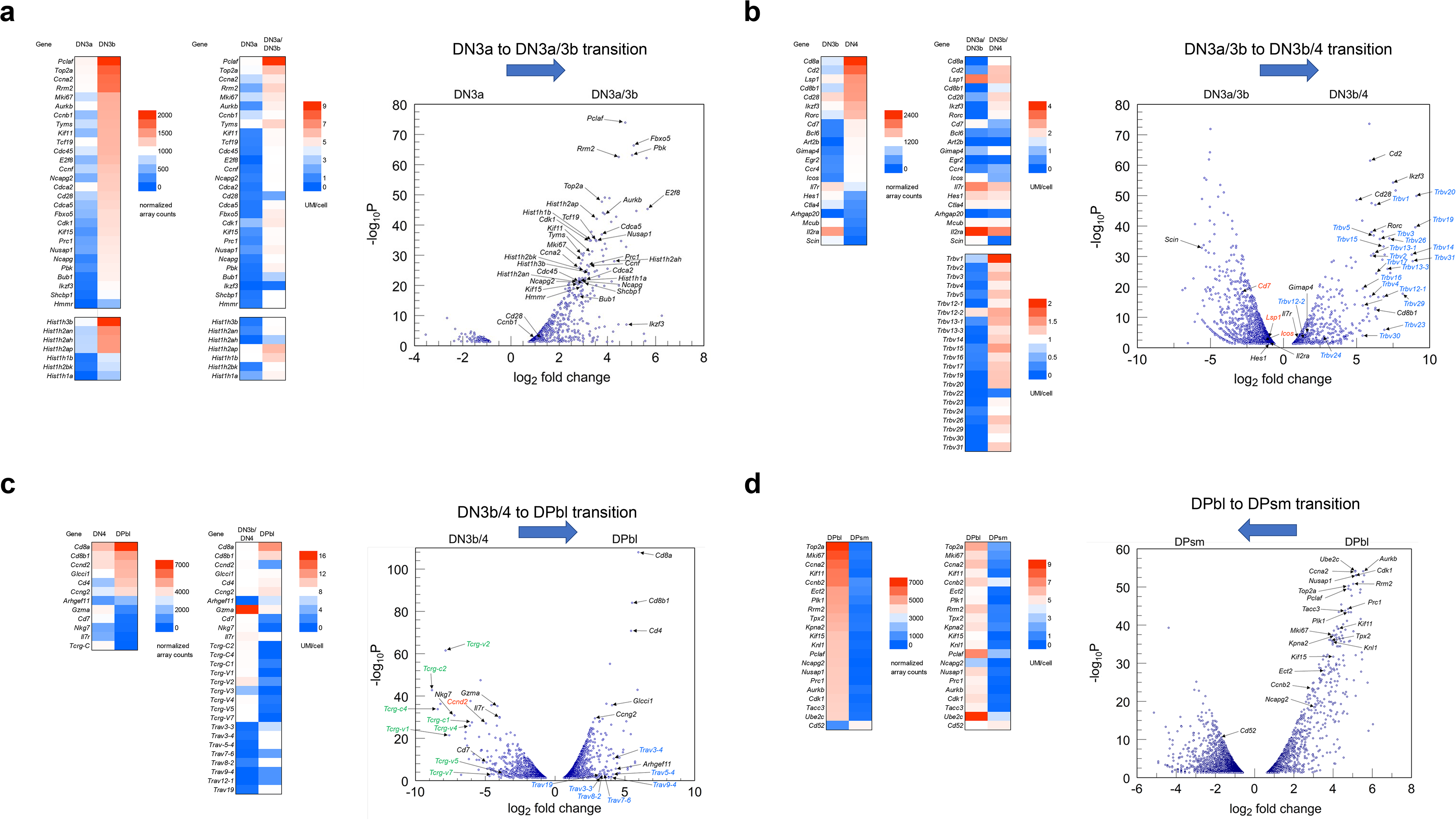
Cluster delineation of DN3a to DPsm cell transitions. For each transition, data from the Immune Genome Project (IGP) microarray and RNA-Seq data was used to construct a panel representing genes with the highest fold-change between phenotypically defined stages of thymocyte differentiation. The gene panel was then used to query the MHC^+^ thymocyte clusters identified by UMAP projection. Combination of library phenotype together with good fit to the interrogating gene panel permitted identification of cluster relationships and developmental trajectories. **a.** Delineation of early post-β selection checkpoint DN3a/3b thymocytes from pre-β selection checkpoint DN3a thymocytes by differential gene expression. The left-hand heatmap depicts a panel selected by comparison of DN3b thymocyte gene expression from the IGP with DN3a cell expression. The same genes were examined for expression in the clusters defined as DN3a and DN3a/3b in Fig.1B (right-hand heatmap). The volcano plot depicts the log_2_-fold increase of expression in the DN3a/3b population over DN3a for the expected normal developmental trajectory (x-axis). Note that for all volcano plots reported here, only the significantly changed transcripts are depicted (P*_adj_* < 0.05; y-axis). **b.** Delineation of late post-β selection checkpoint DN3b/4 thymocytes from early pre-β selection checkpoint DN3a/3b thymocytes by differential gene expression. The heatmap on the far left depicts a panel selected by comparison of DN4 thymocyte gene expression from the IGP with DN3b cell expression (neither DN3a/3b nor DN3b/4 transitional states are explicitly defined in the IGP database). Transcripts in red were predicted from IGP data to be upregulated in the DN3b to DN4 transition but are downregulated for the conditions reported here. **c.** Delineation of late post-β selection checkpoint DN3b/4 thymocytes from DPbl thymocytes by differential gene expression. The DPbl cluster was extracted from the DP library and delineated from the more mature DPsm population by transcriptome signature as described below. **d.** Delineation of mature DPsm thymocytes from cycling DPbl thymocytes by differential gene expression. The heatmap on the far left depicts a panel selected by comparison of DPsm thymocyte gene expression from the IGP with DPbl cell. Note that during the DPbl to DPsm transition, significant cell cycling transcripts were downregulated thus significantly upregulated transcripts in the volcano plot represent the DPbl cells.

**Extended Data Fig. 3.**
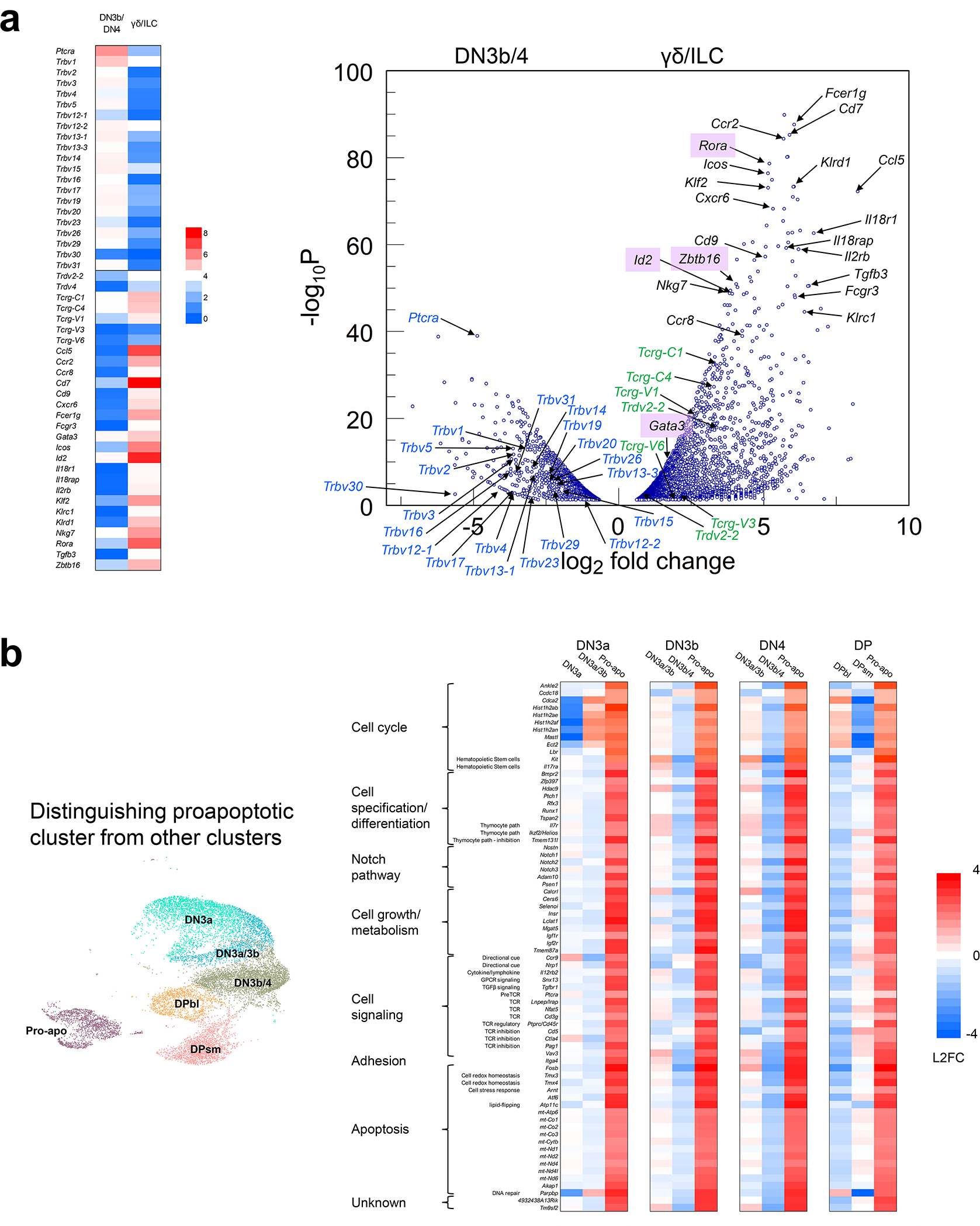
Delineating the ILC-γ/δ TCR thymocyte cluster and pro-apoptotic cluster from the main α/β TCR lineage pathway. **a.** Distinguishing ILC-γ/δ-like cells from DN3b/4 in the DN4 libraries by gene expression. The heatmap on the left shows a manually curated panel of gene transcripts selected by likely high representation in either DN3b/4 or ILC-γ/δ-like cells. Log2 Fold-change (L2FC) and P*_adj_* in the DN4 libraries for differential expression between the DN3b/4 clusters and ILC-γ/δ-like clusters are shown in the volcano plot to the right with transcripts associated with ILC development are highlighted in light purple (*Id2*, *Zbtb16*, *Gata3*, *Rora*). TCR γ and δ transcripts are highlighted in green, and *Trbv* transcripts highlighted in blue. **b**. Gene expression profile of the pro-apoptotic cluster. The dominant pro-apoptotic cluster upregulated gene expression changes are similar between all the MHC^+^ libraries on comparison with the 2 dominant clusters within each of these libraries. All log2-fold changes (L2FC) are relative only to the 3 clusters listed in each heatmap (*i.e.* local) and not to the average across all clusters in that library.

**Extended Data Fig. 4.**
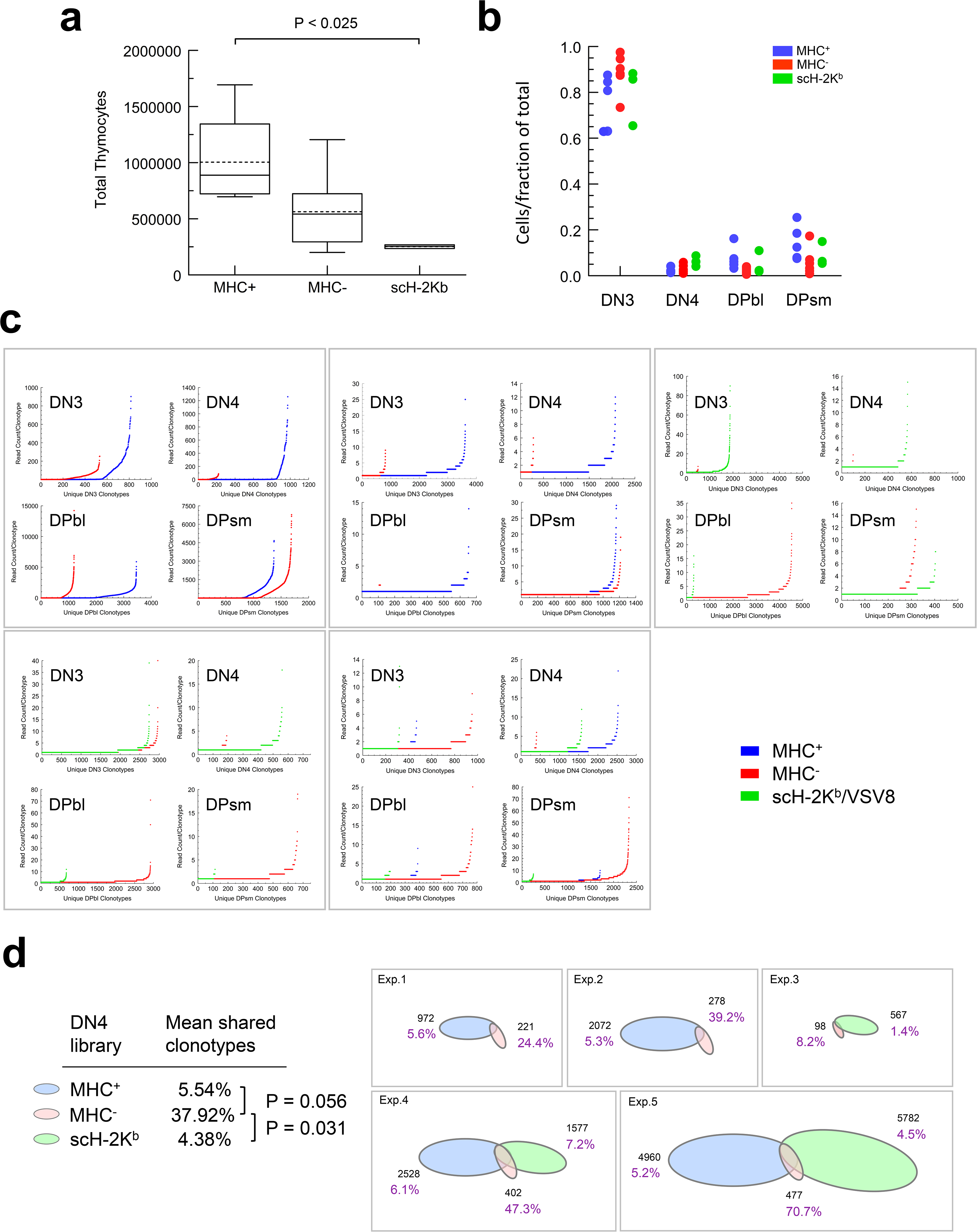
Development and TCR repertoire analyses for cells growing on MHC^+^, MHC^-^ and scH-2K^b^ stromal support cells. **a.** Total cell recoveries after 9d development from 2,000 seeded HSC (6 independent experiments examining 5 independent MHC^+^ cultures, 6 independent MHC^-^ cultures, and 3 independent scH-2K^b^ cultures). Boxes bound 25^th^ to 75^th^ percentile; median is solid line; mean is dotted line; whisker is 1.5 interquartile range (IQR). P determined by t test. **b.** Apparent thymocyte developmental stage representation as fraction of total cells for cultures represented in panel a. **c.** Stage-specific analysis of β chain clonotype representation/10,000 cells in d9 MHC^+^, MHC^-^, and scH-2K^b^ OP9-DL4 development cultures. Representation of data from replicate experiments of data in Fig. 2c-f. **d.** TCR β chain clonotype diversity at DN4 on MHC^+^, MHC^-^, and scH-2K^b^ stroma. The total number of TCR β chain clonotypes (black) recovered from 10^4^ cells of each DN4 population isolated after growth for 9d on the varying OP9-DL4 stroma is represented by an ellipse of area in direct proportion to unique clonotype count (5 independent experiments). Percentage shared clonotypes of the total for each condition (MHC^+^ in blue, MHC^-^ in pink, and scH-2K^b^ in green) is depicted. Note that the area of overlap only approximates degree of sharing to maintain consistent orientation of the ellipses for presentation. The overlap of MHC^+^ and scH-2K^b^ for experiments 4 and 5 is <1% and too small to represent in this format. Statistics and P calculated from Student’s t test presented on left.

**Extended Data Fig. 5.**
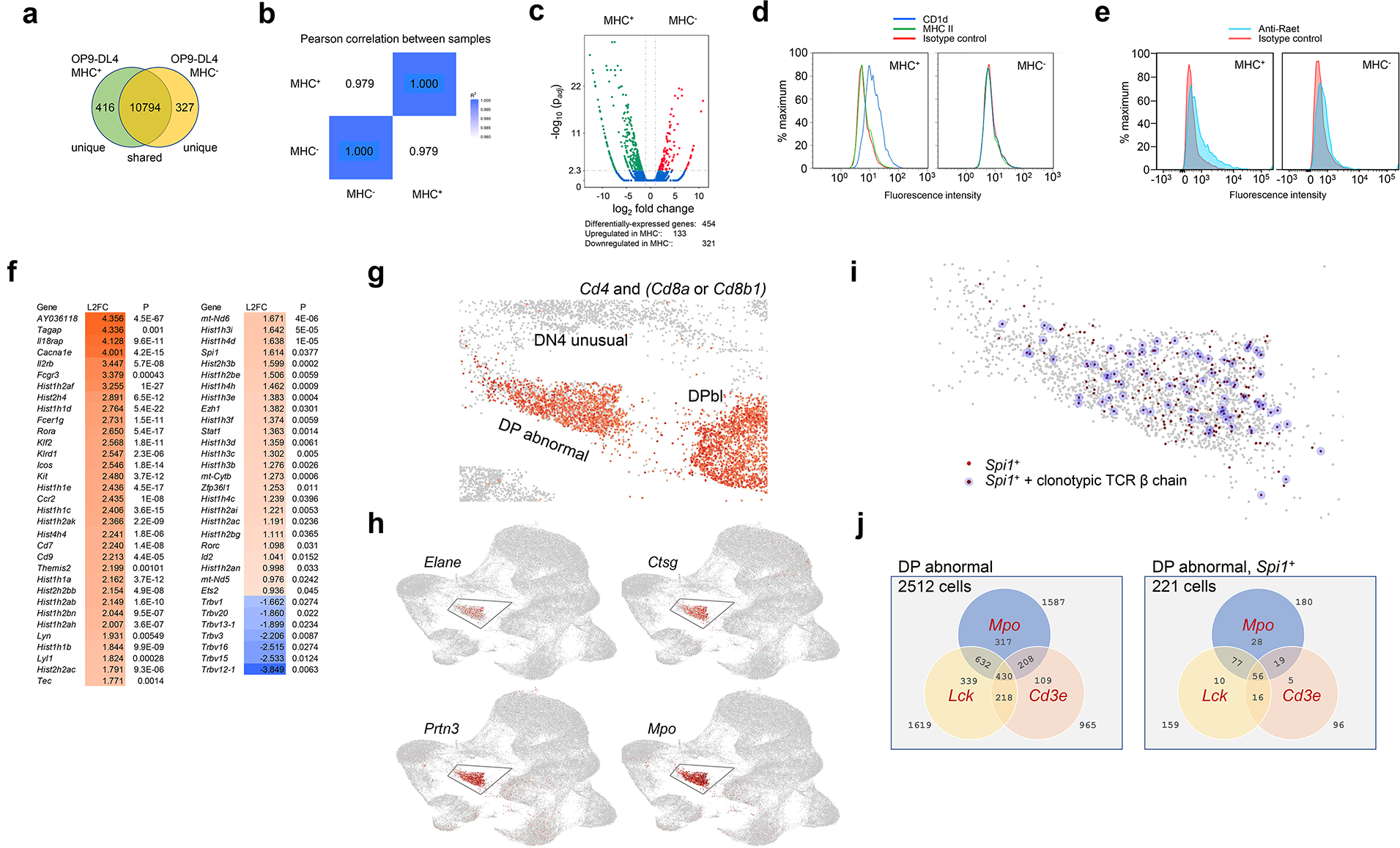
Transcriptome and selected phenotype comparison of MHC^+^ and MHC^-^ OP9-DL4 cells and select gene expression profiles for the DN4 unusual and DPbl abnormal populations. **a.** Comparison of MHC^+^ and MHC^-^ OP9-DL4 stromal cells for transcriptome and phenotypic differences. 93.6% of transcripts detected shared by MHC^+^ and MHC^-^ stroma. **b**. Correlation between cell transcriptomes. Square of Pearson correlation coefficient (R^2^ = 0.958) ideally greater than 0.92 under optimal experimental conditions. **c**. Differential gene expression is <4% of all transcripts detected. **d**. Loss of CD1d surface expression in *B2m/Tap2* KO MHC^-^ OP9-DL4 and confirmation of lack of MHC Class II expression in MHC^+^ and MHC^-^ OP9-DL4. **e**. *Raet* expression in MHC^+^ and MHC^-^ OP9-DL4. **f.** Select transcripts significantly differentially expressed between the MHC^-^ DN3b/4 cluster and the DN4 “unusual” cluster. Heatmap depicts log_2_-fold change (L2FC) of the DN4 “unusual” cluster relative to the DN3b/4 cluster. Actual L2FC values are listed within the heatmap. **g.** Co-expression of *Cd4* transcript with *Cd8a* and/or *Cd8b1* transcripts in an overlay of the MHC^-^ libraries focussed on the DN4 unusual, DPbl, and DP abnormal clusters. **h.** Characteristic myeloid gene transcript expression maps to the MHC^-^ DP abnormal cluster. **i.** Full-length clonotypic TCR β chain transcript expression in 82 of 221 *Spi1*^+^ cells (37.1%) in the MHC^-^ DP abnormal cluster. **j.** *Mpo*-expressing cells in the DP abnormal cluster and the *Mpo*^+^*Spi1*^+^ subset co-express T lineage *Lck* and/or *Cd3e*.

**Extended Data Fig. 6.**
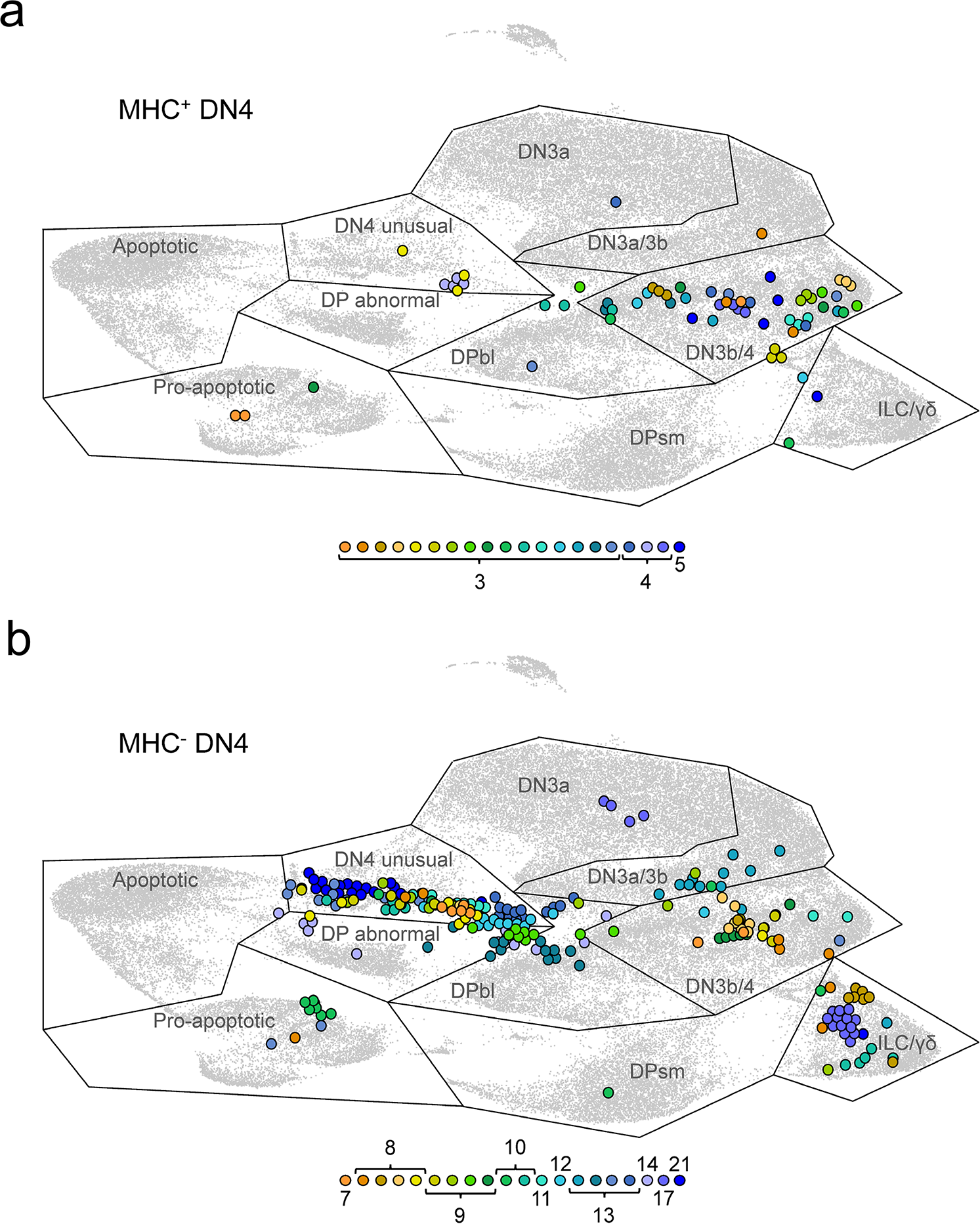
Highly proliferating clonotypic progeny cluster together by transcriptional signature. a. MHC^+^ DN4 20 most highly represented clonotypes by cell number. b. MHC^-^ DN4 20 most highly represented clonotypes by cell number. The identical MHC^+^ and MHC^-^ DN4 clonotypic cells to those presented in Fig. 3d and Extended Data Table 3a are shown in their mapped positions in the UMAP projection. Each clonotype is represented for each panel in a unique colour with cell number indicated in key. Note that colours are not directly related to those used in Fig. 3d.

**Extended Data Fig. 7.**
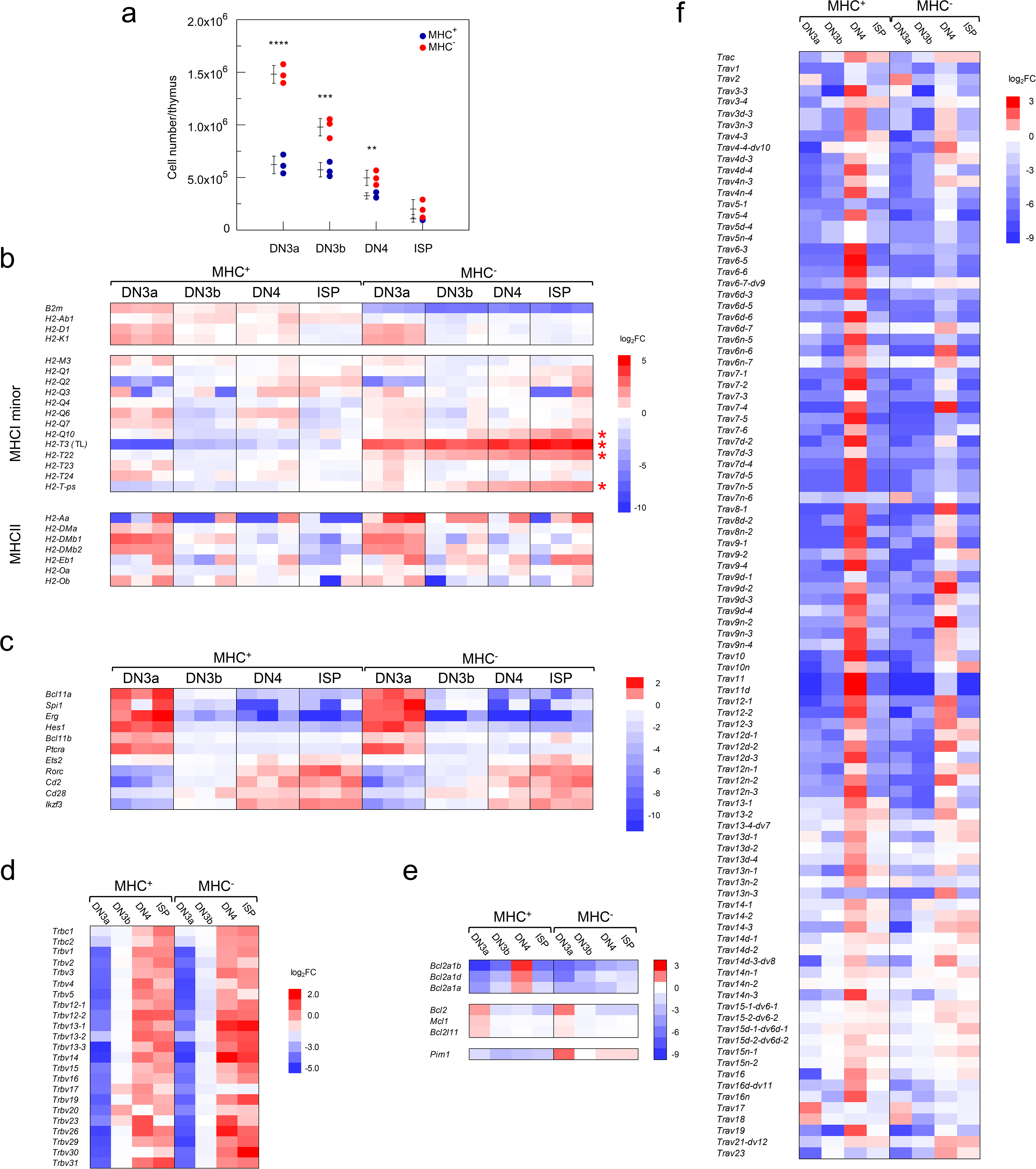
Transcriptome comparison of DN and ISP thymocyte subsets from MHC^+^ and MHC^-^ mice. a. Thymocyte subset cell recoveries from thymi of MHC^+^ and MHC^-^ mice. Mean ± S.D. shown; 3 mice/group; ** P < 0.02; *** P < 0.005; **** P < 0.0005. b. Log_2_-fold change in expression from global population mean for the MHC^-^ knocked out genes (*B2m*, *H2-Ab1*), classical and minor MHCI genes, and MHCII genes. Note that for each thymocyte subset there are 3 replicates except for the MHC^-^ DN4 cells for which there are duplicates. Asterisks highlight transcripts that are upregulated across all MHC^-^ libraries on comparison with MHC^+^ *Q10* (P = 7 x 10^-5^), *H2-T3 (TL)* (P = 3 x 10^-7^), *H2-T22* (P = 1 x 10^-7^) and *H2-T-ps* (P = 4 x 10^-5^). c. Log_2_-fold change in expression of all development stage marker genes depicted in Fig. 1a. d. Log_2_-fold change in TCR Vβ chain segment (*Trbv*) expression. Mean depicted of triplicates for all libraries except for duplicates for MHC^-^ DN4 samples. e. Log_2_-fold change in *Bcl2a1* family transcripts (upper panel), canonical *Bcl2* transcripts (middle panel), and *Pim1* protooncogene (lower panel). Mean values presented. f. Log_2_-fold change in TCR Vα chain segment (*Trav*) expression. Mean depicted of triplicates for all libraries except for duplicates for MHC^-^ DN4 samples.

**Extended Data Table 1.**
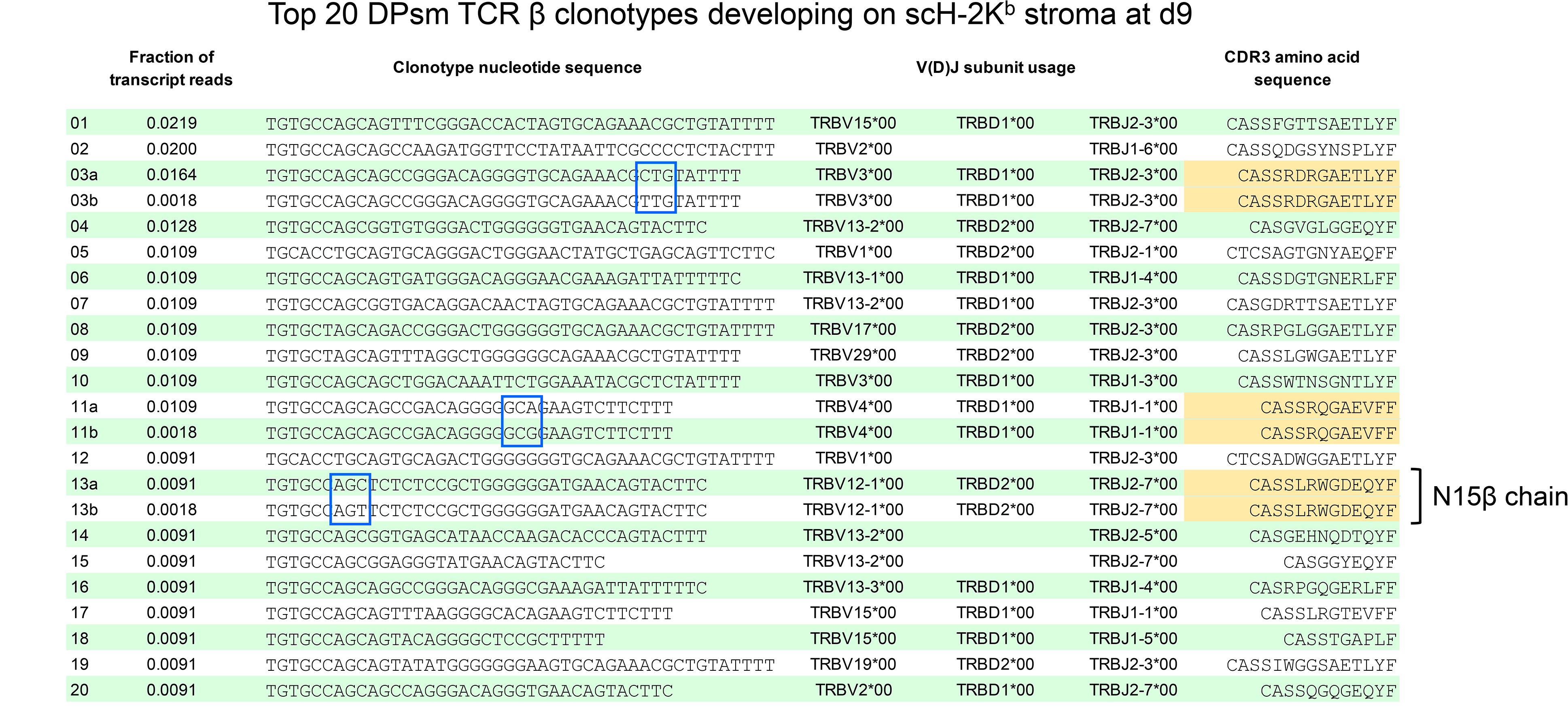
Top 20 DPsm TCR β clonotypes developing on scH-2K^b^ stroma at d9. Codons highlighted by blue border indicate unique clonotypes encoding the same CDR3 amino acid sequence (in gold). The N15β chain is highlighted with known specificity for interaction with VSV8 peptide presented by H-2K^b^. This interaction may occur in the context of a TCR α chain (αβTCR at the DP stage) or absence of a mature TCR α chain (preTCR at the DN3 stage).

**Extended Data Table 2.**
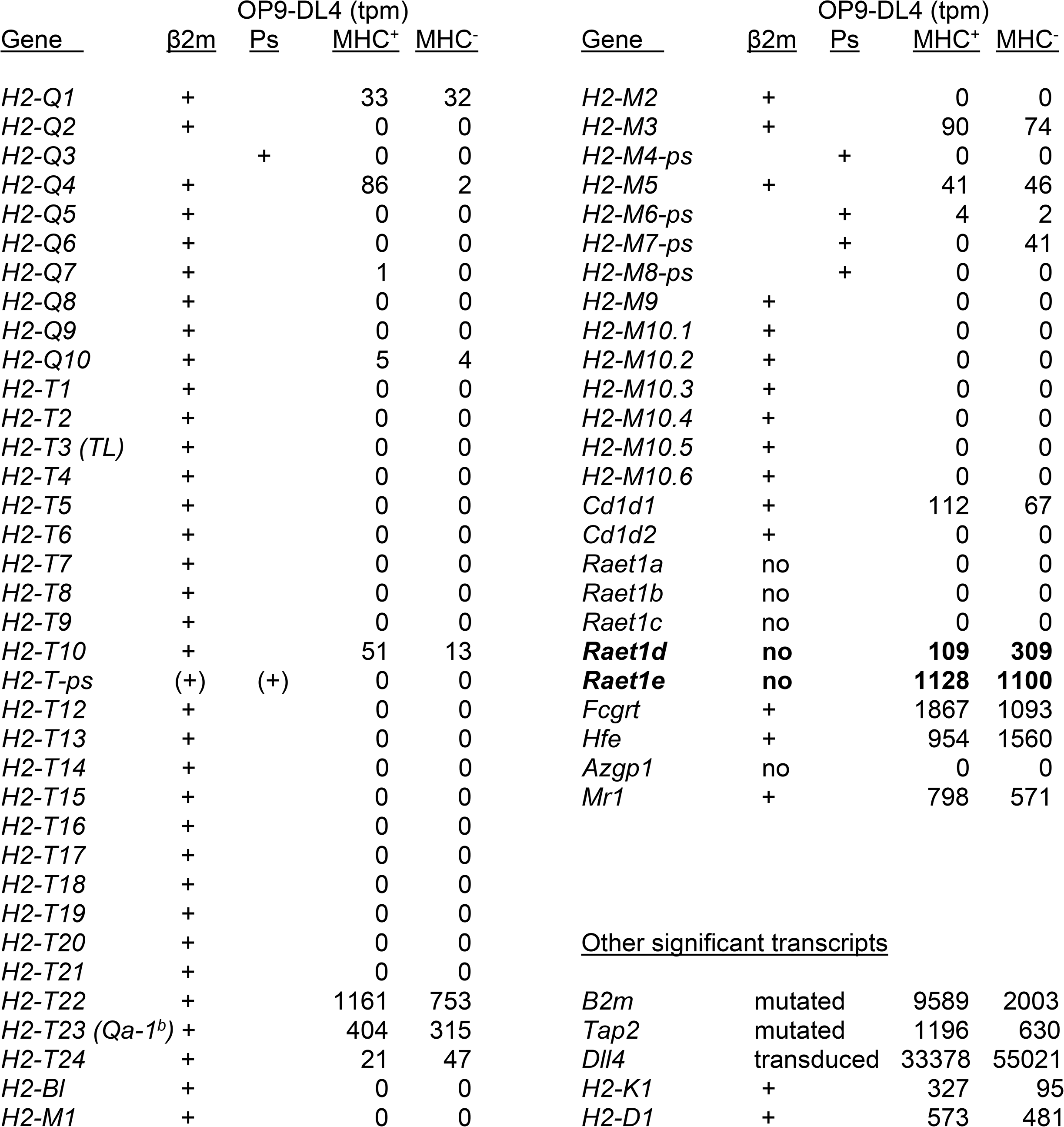
Non-classical MHC class I expression in MHC^+^ and MHC^-^ OP9-DL4. Sixty-one non-classical MHC are listed^66^. Dependence on β2m indicated by ‘+’ or ‘no’. Genes labeled as pseudogene (Ps) may result in transcripts, initially classified as non-coding but subsequently found to be protein coding, as in the instances of *H2-Q5*, *H2-Q10, H2-T1, H2-T4, H2-T12, H2-T13, H2-T14, H2-M10.4, H2-M10.6*. *H2-T-Ps* has been provisionally redefined as protein coding. Genes in bold font are not β2m-dependent and have transcript levels above zero measured as transcripts per million (tpm). Small panel at bottom right presents data for the CRISPR/Cas9 targets *B2m* and *Tap2* deleted in the MHC^-^ variant, Delta-like ligand 4, and major MHCI alleles.

**Extended Data Table 3.**
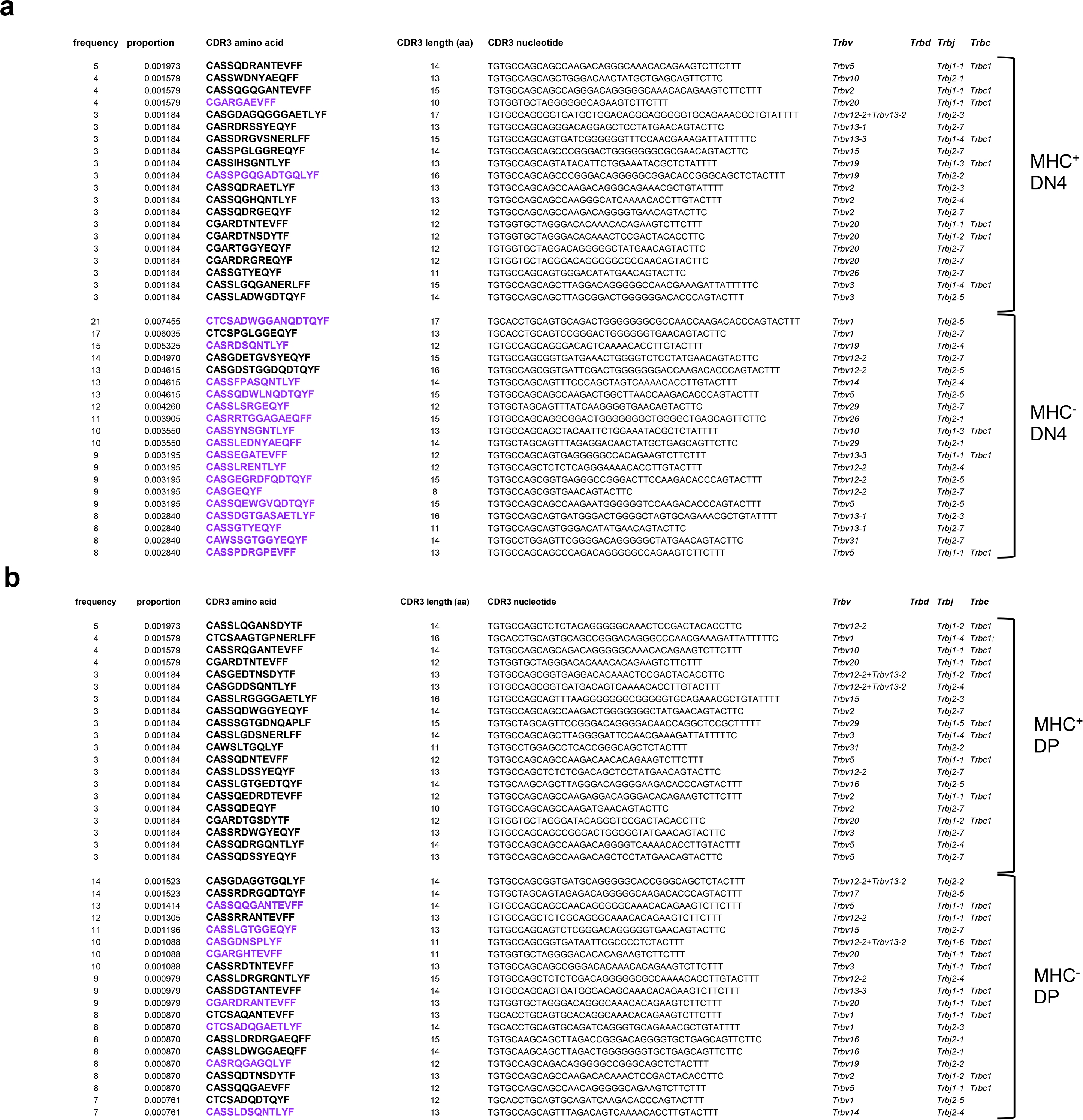
Well-represented clonotypes in MHC^+^ and MHC^-^ libraries. **a.** Top 20 clonotypes in the DN4 libraries; clonotypes present in the DN4 unusual cluster are highlighted in purple. Note that the *Trbv12-2+Trbv13-2* transcript is not a mix of *Trbv12-2* and *Trbv13-2* but rather the result of an independent recombination event between the 5’ end of *Trbv12-2* and the 3’ end of *Trbv13-2*, an event recently and frequently detected in 10X TCR single cell repertoire analyses. **b.** Top 20 clonotypes in the DPbl libraries; clonotypes present in DP abnormal cluster are highlighted in purple.

## Supplementary Information

**File 1.** Gene expression level (UMI/cell), log_2_fold change (global), and P*_adj_* for all clusters in all MHC^+^ libraries – scRNA-Seq.

MS Excel file: Supplementary-Information-File1.xlsx

**File 2.** Gene expression level (UMI/cell), log_2_fold change (global), and P*_adj_* for all MHC^-^ clusters – scRNA-Seq.

MS Excel file: Supplementary-Information-File2.xlsx

**File 3.** TCR β chain clonotypes for DN3, DN4, DPbl and DPsm thymocytes developing on MHC^+^, MHC^-^, and scH-2K^b^/VSV8 stroma.

MS Excel file: Supplementary-Information-File3.xlsx

**File 4.** Representative FACS separation profiles of developing thymocyte-like subsets *in vitro* and thymocyte subsets *in vivo*.

PDF file: Supplementary-Information-File4.pdf

**File 5.** Gene expression and total TCRβ clonotype repertoire data for DN3a, DN3b, DN4 and ISP cells isolated from MHC^+^ and MHC^-^ mice.

MS Excel file: Supplementary-Information-File5.xlsx

**File 6.** Gene expression analysis of a chromosome Y transcript panel and expression-matched autosomal panel to address aberrant clonal HSC expansion contributing to the MHC^-^ DN4 unusual and DP abnormal populations.

MS Excel file: Supplementary-Information-File6.xlsx

